# Brain-predicted age difference score is related to specific cognitive functions: A multi-site replication analysis

**DOI:** 10.1101/652867

**Authors:** R. Boyle, L. Jollans, L.M. Rueda-Delgado, R. Rizzo, G.G. Yener, J.P. McMorrow, S.P. Knight, D. Carey, I.H. Robertson, D.D. Emek-Savaş, Y. Stern, R.A. Kenny, R. Whelan

## Abstract

Brain-predicted age difference scores are calculated by subtracting chronological age from ‘brain’ age, which is estimated using neuroimaging data. Positive scores reflect accelerated ageing and are associated with increased mortality risk and poorer physical function. To date, however, the relationship between brain-predicted age difference scores and specific cognitive functions has not been systematically examined using appropriate statistical methods. First, applying machine learning to 1,359 T1-weighted MRI scans, we predicted the relationship between chronological age and voxel-wise grey matter data. This model was then applied to MRI data from three independent datasets, significantly predicting chronological age in each dataset: Dokuz Eylül University (n=175), the Cognitive Reserve/Reference Ability Neural Network study (n=380), and The Irish Longitudinal Study on Ageing (n=487). Each independent dataset had rich neuropsychological data. Brain-predicted age difference scores were significantly negatively correlated with performance on measures of general cognitive status (two datasets); processing speed, visual attention, and cognitive flexibility (three datasets); visual attention and cognitive flexibility (two datasets); and semantic verbal fluency (two datasets). As such, there is firm evidence of correlations between increased brain-predicted age differences and reduced cognitive function in some domains that are implicated in cognitive ageing.

## Introduction

Longitudinal neuropsychological testing in older adults can be used to detect cognitive decline. However, practice effects can obscure assessment of cognitive ability (Elman et al., 2018), and test performance is affected by subject-level factors such as the individual’s level of comprehension, reading ability, self-efficacy, motivation, fatigue, and fluctuations in concentration (McCaffrey & Westervelt, 1995). In contrast, objective biomarkers are not subject to such biases or patients’ physical limitations (Jollans & Whelan, 2016). An objective biomarker of cognitive ageing would therefore be useful for the timely identification of cognitive decline outside of age-related norms.

Ageing is a process with significant heterogeneity across individuals (McCrory & Kenny, 2018). Consequently, chronological age is not the most accurate marker of an individual’s rate of biological ageing (Sprott, 2010). Ageing biomarkers have been developed that provide additional information about an individual’s health status and life expectancy (Dean & Morgan, 1988). For example, DNA methylation data can estimate epigenetic ageing (‘epigenetic clocks’), reflecting the age of an individual’s tissues or blood cells (Fiorito et al., 2019). Subtracting chronological age from the biological age results in a biologically informative summary score – the predicted age difference – for each individual, which reflects the deviation from typical lifespan trajectories (Richard et al., 2018). This approach has also been applied in neuroimaging, where machine learning can be used to quantify the relationship between structural MRI data and chronological age, in order to estimate an individual’s ‘brain age’. Subtracting chronological age from the estimated ‘brain age’ results in a brain predicted-age difference score (*brainPAD*, also referred to as brain age gap, brainAGE, Brain-Age Score; (Beheshti, Maikusa, & Matsuda, 2018; Franke, Ziegler, Klöppel, & Gaser, 2010; Schnack et al., 2016) which quantifies how a person’s brain health differs from what would be expected for their chronological age.

BrainPAD is a promising biomarker of general brain ageing as it already satisfies several criteria for ageing biomarkers (Butler et al., 2004). BrainPAD is predictive of mortality (Cole, Ritchie, et al., 2018) and of age-sensitive physiological measures, including grip strength, lung function, walking speed and allostatic load (Cole, Ritchie, et al., 2018). Moreover, BrainPAD could potentially be used as a biomarker of cognitive ageing as it is negatively correlated with fluid cognitive performance (Cole, Ritchie, et al., 2018) and is significantly increased in Alzheimer’s disease (AD) and mild cognitive impairment (MCI) (Franke & Gaser, 2012; Gaser et al., 2013; Löwe, Gaser, & Franke, 2016). However, the potential of brainPAD as a cognitive ageing biomarker is currently limited by a lack of knowledge regarding the exact relationship between brainPAD and specific cognitive functions in healthy individuals.

Studies relating specific cognitive functions and brainPAD have been assessed in solely clinical samples (e.g., Cole et al. (2015), traumatic brain injury), or in mixed samples of clinical groups and healthy controls (e.g., Beheshti et al. (2018); AD, MCI, and healthy controls) and not samples comprised only of healthy adults. As such, the reported associations between brainPAD and specific domains of cognitive function in such studies (Beheshti et al., 2018; Cole et al., 2015) may be skewed towards significance by the inclusion of the clinical samples with typically higher brainPADs. Consequently, these findings may not represent the brainPAD-cognition relationship in normal ageing. For example, Le and colleagues (2018) reported a significant negative correlation between brainPAD and response inhibition and selective attention in a sample of individuals comprised of healthy controls and patients with mood or anxiety disorders, substance use disorder and/or eating disorders. However, significantly increased brainPADs have been reported in mood disorders such as major depression (Koutsouleris et al., 2014) and in substance use disorders such as alcohol dependence (Guggenmos et al., 2017). As both major depression and alcohol dependence are associated with cognitive impairments (Chanraud et al., 2007; McIntyre et al., 2013), the significant brainPAD-cognitive function correlations reported across samples including such populations could be driven by the inclusion of such clinical groups.

The relationship between specific cognitive functions and BrainPAD has also been somewhat obscured by statistical considerations. Recent work has empirically demonstrated that chronological age must be controlled for when testing relationships between brainPAD and cognitive functions (Le et al., 2018; Smith, Vidaurre, Alfaro-Almagro, Nichols, & Miller, 2019). Failure to correct for chronological age can result in false positive findings because some cognitive variables are correlated with chronological age – but *not* brain ageing – and brainPAD is typically correlated with chronological age (Le et al., 2018). In light of this recent work, it is difficult to interpret studies that did not control for chronological age when investigating the brainPAD-cognition relationship in healthy controls (Franke, Gaser, Manor, & Novak, 2013; Löwe et al., 2016). A second statistical issue is a failure to correct for multiple comparisons. Researchers testing the brainPAD-cognition relationship have tended to carry out multiple statistical tests of the correlation between brainPAD and various cognitive measures. The performance of multiple statistical tests can increase the Type I error and result in false positive findings (Ranganathan, Pramesh, & Buyse, 2016). However, some papers did not control for multiple comparisons when investigating the brainPAD-cognition relationship (Beheshti et al., 2018; Cole, Underwood, et al., 2017). Other studies have investigated the relationship between brainPAD and specific domains of cognitive function while controlling for chronological age and multiple comparisons, but there are conflicting results for most cognitive domains. For example, a significant correlation between verbal fluency and brainPAD was reported by Franke and colleagues (2013) whereas Richard and colleagues (2018) found no association between verbal fluency and brainPAD. We have summarized the brainPAD-cognition findings in Table 1.

**Table 1.**
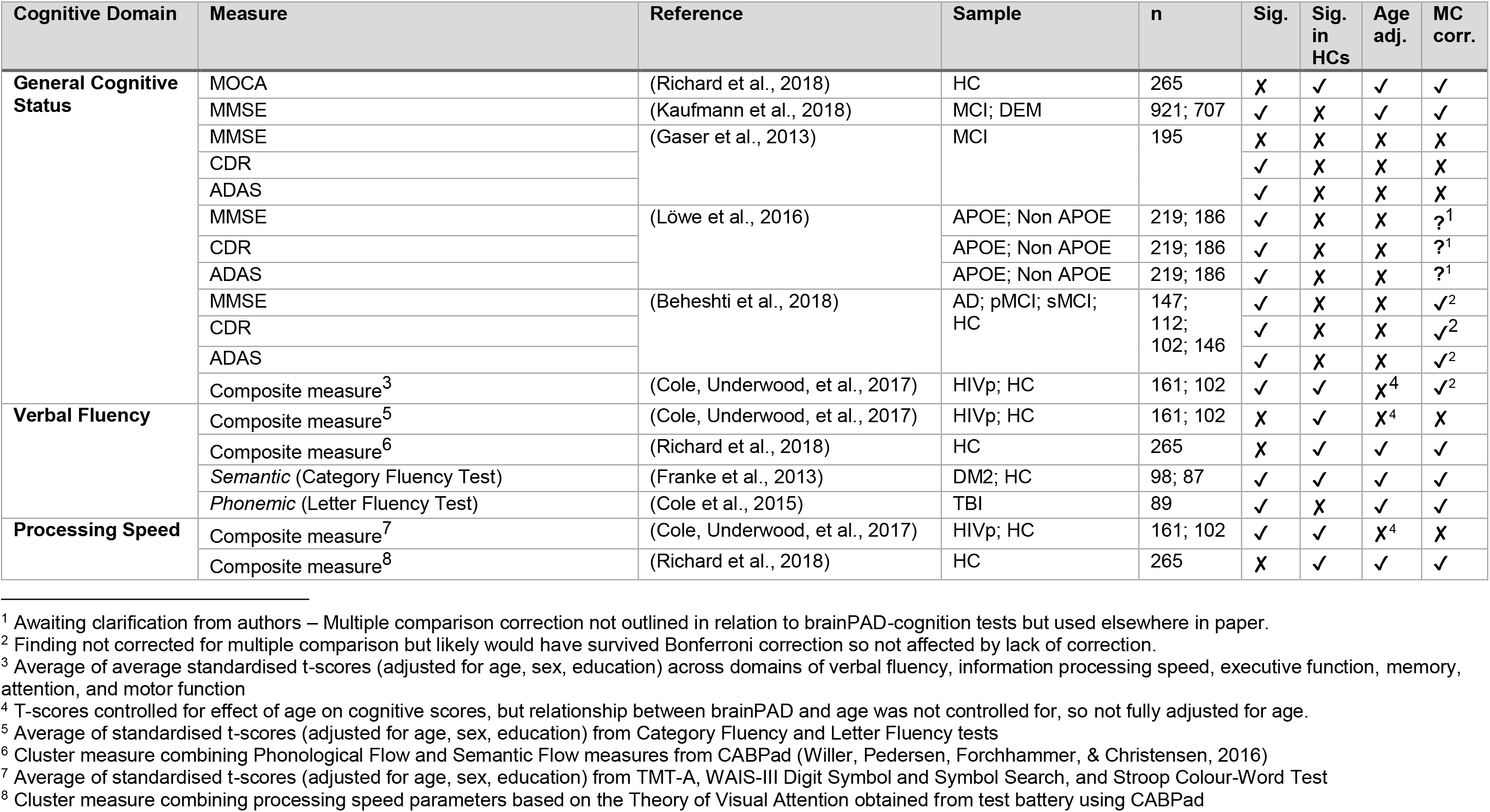
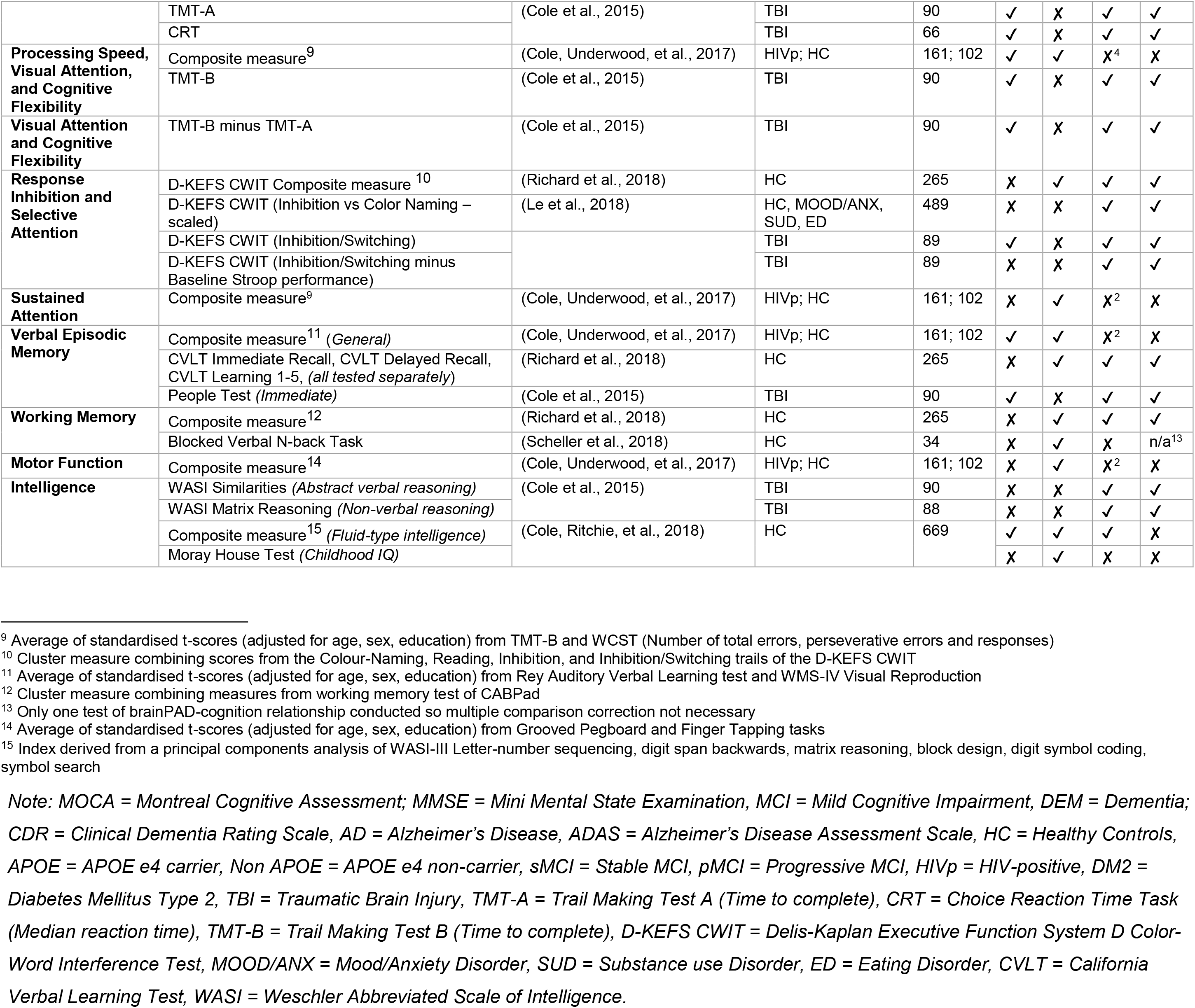
Summary of findings on the relationship between brainPAD and cognitive function, indicating whether results were statistically significant (Sig.), statistically significant in healthy controls (Sig. in HCs), adjusted for age (Age adj.) and corrected for multiple comparisons (MC corr.).

The first step in generating a brainPAD score is creating a feature set of neuroimaging data which is correlated with chronological age. Neuroimaging data have high dimensionality, which can result in overfitting and overoptimistic predictions (Whelan & Garavan, 2014). Brain age prediction models thus rely on feature engineering techniques such as principal components analysis (PCA; Franke et al., 2010; Gutierrez Becker, Klein, & Wachinger, 2018) or even dot products of different features (e.g. vectors of GM and white matter (WM) voxels as in Cole et al., 2015; Cole, Ritchie, et al., 2018; Cole, Underwood, et al., 2017) in order to reduce the dimensionality (Mwangi, Tian, & Soares, 2014). Although these models ultimately create generalizable and accurate predictions, these come at the cost of reduced interpretability of the contributions of the features (Bunea et al., 2011; Mateos-Pérez et al., 2018), which is important for assessing the neurobiological validity of the model (Woo, Chang, Lindquist, & Wager, 2017) and to identify specific brain areas for further investigation (Scheinost et al., 2019). An alternative to applying PCA or other data reduction techniques is to use penalized regression methods such as the Elastic Net (Zou & Hastie, 2005), with only one class of features (e.g. GM voxels) as input. GM data is particularly well-suited for age prediction as GM volume linearly declines with age (but cf. Fjell et al., 2013) whereas WM volume has a less straightforward relationship with age, as it doesn’t decline significantly until middle age (Farokhian, Yang, Beheshti, Matsuda, & Wu, 2017; Ge et al., 2002). The Elastic Net is a machine learning model well-suited to the high dimensionality and multicollinearity inherent in neuroimaging data as shown by the finding that it produced the most consistent predictions as compared to various other models over datasets with varying sample-, feature set-, and effect-sizes (Jollans et al., in revision).

A final challenge in the development of neuroimaging biomarkers, or neuromarkers, is ensuring the generalisability of the neuromarker to new data. For practical reasons, cross-validation, where a dataset is split into a training set and a test set (Varoquaux et al., 2017), is often used as an estimate of model accuracy for new data (Jollans & Whelan, 2018; Scheinost et al., 2019). However, cross-validation accuracy estimates are often optimistically biased and can vary considerably (Varoquaux et al., 2017), particularly when preprocessing and feature selection are carried out on the entire dataset before splitting it into training and test sets (Dwyer, Falkai, & Koutsouleris, 2018; Woo et al., 2017). As such, the gold-standard for assessing the external validity and generalisability of a neuromarker is by testing how the model performs on a completely independent held-out dataset (Jollans & Whelan, 2018). While various brainPAD studies have externally validated their models (Beheshti et al., 2018; Cole et al., 2015; Cole, Ritchie, et al., 2018; Cole, Underwood, et al., 2017; Franke et al., 2010; Gutierrez Becker et al., 2018; Lancaster, Lorenz, Leech, & Cole, 2018; Liem et al., 2017; Madan & Kensinger, 2018; Varikuti et al., 2018), only a few studies have reported model performance in terms of accuracy (i.e., correlation or mean absolute error between brain-predicted age and chronological age) on the external validation dataset (Cole et al., 2015; Lancaster et al., 2018; Liem et al., 2017; Madan & Kensinger, 2018). This does not necessarily cast doubt on the validity of the models whose accuracy is reported in terms of internal cross-validation performance. However, not reporting the external validation performance limits the interpretation of the accuracy and generalisability of various brainPAD models as typically performance will be lower in the external validation dataset.

In order to clarify the unclear relationship between brainPAD and specific domains of cognitive function, we aimed to 1) establish an interpretable model of brainPAD using the Elastic Net with GM voxel-wise data, 2) externally validate this model in three independent datasets, and 3) to establish the domains of cognitive function that are reliably correlated with brainPAD across different datasets.

## Methods

### Participants

#### Training Set

The data were comprised of MRI scans from 1,359 healthy adults (mean age 40.04 years, SD = 17.78 years, range = 18.00 - 88.36 years; 855 females) drawn from various open-access data repositories (see Table S.1 in Supplementary Info). Inclusion criteria for the training cohort were: over 18 years old, age and gender data available, and not diagnosed with any neurological, psychiatric or major medical conditions.

#### Independent Test Set 1 – Dokuz Eylül University (DEU)

The first test set was comprised of 175 community-dwelling adults (mean age = 68.95 years, SD = 8.59 years; range = 47.56 – 93.51 years; 104 females) recruited as part of a study conducted at Dokuz Eylül University, Izmir, Turkey. Exclusion criteria included history of neurological or psychiatric diseases, use of psychotropic drugs including cholinesterase inhibitors, traumatic brain injury, history of stroke, drug and/or alcohol addiction and uncontrolled systemic diseases.

#### Independent Test Set 2 – Cognitive Reserve/Reference Ability Neural Network (CR/RANN)

The third test set was comprised of 380 community-dwelling adults (mean age = 52.41 years, SD = 17.09 years; range = 19 – 80 years; 210 females) who participated in the Cognitive Reserve/Reference Ability Neural Network study (CR/RANN; (Stern, Gazes, Razlighi, Steffener, & Habeck, 2018; Stern et al., 2014). These participants were screened for MRI contraindications, hearing and visual impairments, medical or psychiatric conditions, and dementia and MCI. Further inclusion criteria were a score of over 135 on the Mattis Dementia Rating Scale (Jurica, Leitten, & Mattis, 2001), a reading level at least equivalent to the US 4^th^ grade, and minimal complaints of functional impairment.

#### Independent Test Set 3 – The Irish Longitudinal Study on Ageing (TILDA)

The second test set was comprised of an MRI subset of a nationally representative longitudinal cohort study of community-dwelling adults in Ireland (B. J. Whelan & Savva, 2013). From an initial subset of 502 participants, participants were excluded due to missing a portion of the cerebellum (n = 2), a history of Parkinson’s disease, stroke, or transient ischemic attack (n = 11) and no cognitive data (n= 2). The final test set was comprised of MRI data from 487 participants (mean age = 68.6 years, SD = 7.21 years; range = 50 – 88 years; 260 females).

### MRI data acquisition

#### Training Set

A range of T1-weighted MRI scans from different scanners and using different protocols were used as the training set (see Table S.1 in Supplementary Info).

#### Test Set 1 – DEU Dataset

DEU participants underwent a 10 minute T1 scan in a 1.5 T Philips Achieva scanner as part of a larger 20-min MRI battery. Two separate protocols were used for scans included here. The Alzheimer’s Disease Neuroimaging Initiative (ADNI) T1 protocol was followed for 126 scans using the turbo field echo sequence with the following parameters: number of slices = 166, FOV = 240mm^3^, slice thickness = 1 mm, slice gap = 0 mm, TR = 9 ms, TE = 4 ms. For 49 scans, a local protocol using a gradient echo sequence was followed with the following parameters: FOV = 230mm^3^, slice thickness = 1 mm, slice gap = 0 mm, TR = 25 ms, TE = 6 ms.

#### Test Set 2 – CR/RANN Dataset

CR/RANN participants underwent a 5 minute T1 MPRAGE scan in a 3T Philips Achieva scanner as part of a larger 2-hr imaging battery. The following parameters were used: FOV = 256×256 mm, slice thickness = 1 mm, slice gap = 0 mm, TR = 6.5 ms, TE = 3 ms.

#### Test Set 3 – TILDA Dataset

TILDA participants underwent a 5 minute 24 seconds T1 MPRAGE scan in a 3T Philips Achieva scanner as part of a larger 45-min MRI battery. The following parameters were used: FOV = 240×218×162mm^3^, slice thickness = 0.9 mm, slice gap = 0 mm, TR = 6.7 ms, TE = 3.1 ms.

### MRI pre-processing

All images were preprocessed using SPM12 (University College London, London, UK). Prior to processing, all scans were automatically approximately reoriented (see Supplemental Information; MRI pre-processing) to a canonical SPM template. All scans were then visually inspected for good orientation and gross artefacts before preprocessing. In the test set, badly oriented scans were manually reoriented before preprocessing. In both training and test sets, each individual dataset was preprocessed in a separate batch. Bias correction was applied to image which were then segmented into GM, WM, and CSF. Segmented GM images were non-linearly registered to a custom template, using SPM’s *DARTEL*. Images were then affine registered to MNI space (1 mm^3^) and resampled with modulation to preserve the total amount of signal from each voxel. Images were smoothed with a 4 mm full-width at half maximum Gaussian kernel. Finally, images were visually inspected for accurate segmentation. The code used to auto-reorient and preprocess the MRI data is available at https://github.com/rorytboyle/brainPAD.

### Machine learning

#### Data preparation

GM images were resized to 2 mm^3^ voxels and individual voxel values were extracted from each image. In the training set, voxels with a GM density of > 0.2 in all 1,359 images were utilized in the training set. The training data consisted of 1,359 images, each with 54,869 voxels.

#### Machine learning model

The goal of the training phase was to construct a generalizable model that could predict chronological age from GM data. In order to increase generalizability, a data resampling ensemble approach was used. That is, 500 participants, with a 50:50 gender ratio, were randomly sampled without replacement from the training data to form a nested training set. This process was repeated 25 times, creating 25 nested training sets. Each nested training set (500 participants x 54,869 voxels), was used as the input to a regularized linear regression model (Elastic Net), with 10-fold cross-validation (CV), to predict the chronological age of each participant (see Supplementary Info for further information on the machine learning model). The performance of the model was quantified using the mean of each of the 25 nested models’ Pearson’s correlation between chronological age and predicted age (*r*), total variance explained (R^2^), mean absolute error (MAE), and the weighted MAE. The weighted MAE is equal to the MAE divided by the age range of the sample tested and is a more suitable metric for comparing the MAE of brainPAD models across studies as it accounts for the impact of a sample’s age range on prediction accuracy (Cole, Franke, & Cherbuin, 2018). A lower weighted MAE reflects greater accuracy.

#### Application to independent test sets

First, the average coefficient value for each voxel across all folds in all 25 training models was calculated, resulting in a vector of length 54,869. For each independent test set, the mean coefficient values were multiplied by the voxels’ GM density values and the product was summed to create a brain-age prediction for each participant. To correct for the proportional bias in the model, the prediction was added to the intercept of the training set, and the result was then divided by the slope of the training set. This correction does not affect the relationship between brainPAD and outcome measures but scales the data correctly so that brainPAD scores can be interpreted in units of years proportional to a person’s chronological age. Similar corrections have been applied in other brainPAD models (Cole, Ritchie, et al., 2018). BrainPAD was calculated by subtracting chronological age from the corrected predicted age, hence, a positive brainPAD value indicates a brain-predicted age that exceeds the participant’s chronological age, suggesting accelerated brain ageing. The code used to make brain-age predictions and calculated brainPAD scores for independent test sets is available at https://github.com/rorytboyle/brainPAD.

### Cognitive function measures

Each of the three datasets contained a wide range of cognitive measures. For the purposes of the present study, cognitive measures were selected for analysis if a comparable measure existed in at least two of the three datasets. Across all three datasets, 17 common cognitive domains were identified (see Table 2 for list of cognitive domains and cognitive measures used and Supplementary Information for detailed descriptions of each cognitive measure).

**Table 2.** Cognitive measures available across each dataset in comparable cognitive domains (see Table S.2 for full information on each measure).

| **Cognitive Domain(s)** | **DEU Measure (N)** | **CR/RANN Measure (N)** | **TILDA Measure (N)** |
| --- | --- | --- | --- |
| **General Cognitive Status** | MMSE (total score) (172) | DRS (370) | MMSE (485) |
| **Premorbid Intelligence** | n/a | AMNART (362) | NART (486) |
| **Phonemic Verbal Fluency** | KAS Test (137) | CFL Test (360) | n/a |
| **Semantic Verbal Fluency** | Animals Test (175) | Animals Test (361) | Animals Test (487) |
| **Processing Speed** | TMT A (93) | TMT A (361) | CTT 1 (487) |
| **Processing Speed, Visual Attention, Cognitive Flexibility** | TMT B (84) | TMT B (357) | CTT 2 (482) |
| **Visual Attention, Cognitive Flexibility** | TMT B minus TMT A (84) | TMT B minus TMT A (357) | CTT 2 minus CTT 1 (482) |
| **Cognitive Flexibility** | WCST Perseverative Error Percentage (50) | WCST Perseverative Error Percentage (327) | n/a |
| **Response Inhibition, Selective Attention** | Stroop (Turkish Capa version; Emek-Savaş, Yerlikaya, Yener, & Öktem, 2019) Interference Score - Time (150) | Stroop (Golden version; Golden, 1978) Interference Score - Words (359) | n/a |
| **Sustained Attention (Errors of Commission)** | n/a | PVT False Alarms (176) | SART Errors of Commission (482) |
| **Sustained Attention (Reaction Time)** | n/a | PVT Median Reaction Time (176) | SART Coefficient of Variation in Reaction Time (479) |
| **Verbal Episodic Memory (Immediate)** | OVMPT Immediate Recall (175) | SRT Total Score (360) | Immediate Recall (487) |
| **Verbal Episodic Memory (Delayed)** | OVMPT Delayed Recall (175) | SRT Delayed Recall (360) | Delayed Recall (487) |
| **Verbal Episodic Memory (Learning)** | OVMPT Total Learning Score (175) | SRT Consistent Long Term Retrieval (360) | n/a |
| **Working Memory** | WMS-R Digit Span Forward Test (171) | WAIS-III Letter Number Sequencing Test (360) | n/a |
|  | WMS-R Digit Span Backward Test (170) |  |  |
| **Visuospatial Ability** | BLOT (80) | WAIS-III Block Design Test (356) | n/a |

*Note: MMSE = Mini-mental state examination (Folstein, Folstein, & McHugh, 1975); DRS Total Score = Mattis Dementia Rating Scale-2 – Total Score (Jurica et al., 2001); NART = National Adult Reading Test (Nelson & Willinson, 1982); AMNART = American National Adult Reading Test (Grober & Sliwinski, 1991); CTT = Colour Trails Test (D'Elia, Satz, Uchiyama, & White, 1996); TMT = Trail Making Test (Reitan, 1955); WCST = Wisconsin Card Sorting Test (Heaton, Chelune, Talley, Kay, & Curtiss, 1993); SART = Sustained Attention to Response Test (Robertson, Manly, Andrade, Baddeley, & Yiend, 1997); PVT = Psychomotor Vigilance Task (Dorrian, Rogers, & Dinges, 2005); OVMPT = Öktem Verbal Memory Processes Test (Öktem, 1992); SRT = Selective Reminding Test (Buschke & Fuld, 1974); WMS-R = Wechsler Memory Scale (Wechsler, 1987); WAIS-III = Wechsler Adult Intelligence Scale – Third Edition (Wechsler, 1997); BLOT = Benton's Judgement of Line Orientation Test (Benton, Varney, & Hamsher, 1978)*

### Statistical analysis

The statistical analysis was conducted using the following procedure:

1. <u>*Correlate*</u>. Within each independent test set, partial Spearman’s rank order correlations were conducted between brainPAD scores and cognitive measures, controlling for chronological age and gender. Gender was adjusted for to account for a significant gender difference in brainPADs (p < 0. 0001), see Supplementary Results for further detail.
2. <u>*Replicate*</u>. For findings replicated in multiple datasets, the probability of obtaining p-values by chance was calculated by random-label permutation (see Supplementary Methods for further detail). Briefly, this involved randomly shuffling brainPAD scores, conducting Spearman’s partial correlations between randomly shuffled brainPAD scores and the cognitive dependent variables, controlling for age and gender. This process was repeated one million times. The number of times in which all random p-values were more extreme (i.e. smaller) than the actual p-values was summed and divided by one million to obtain the probability of the finding replicating across multiple datasets by chance. Replicated findings were deemed significant if this probability was less than .05.
3. <u>*Correct for multiple comparisons*</u>. All other correlations were then corrected for multiple comparisons, while allowing for correlations among dependent cognitive variables, using a maximum statistic approach (see Supplementary Methods for further detail). Briefly, in each test set, brainPAD scores were randomly shuffled and then Spearman’s partial correlations were conducted between the randomly shuffled brainPAD scores and the cognitive dependent variables, controlling for age and gender. This process was repeated 10,000 times and the maximum rho value was stored each time. Correlations between actual brainPAD scores and cognitive variables were deemed significant if they exceeded the 95th percentile of the maximum rho values.

## Results

### Brain age prediction

#### Training set

The model accurately predicted chronological age (r = 0.85, R^2^ = 67.24%, MAE = 7.28 years, weighted MAE = 0.10, p < 0.0001). As with other brain PAD models (e.g., Cole et al., 2018), a proportional bias was observed in this model where chronological age correlated with prediction error (r = - 0.4452, p = 1.1036e-10).

#### Independent test sets

**Table 3.**
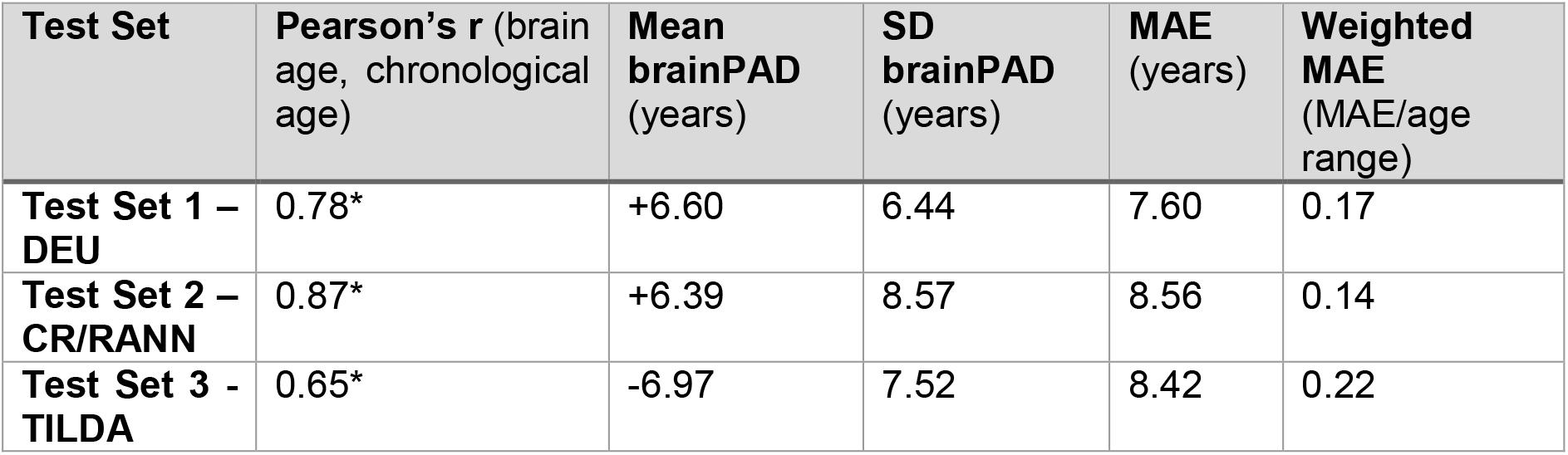
Results of application of trained model parameters to 3 independent test sets. * = *p* < 10^-37^.

| Test Set | Pearson's $r$ (brain age, chronological age) | Mean brainPAD (years) | SD brainPAD (years) | MAE (years) | Weighted MAE (MAE/age range) |
| --- | --- | --- | --- | --- | --- |
| Test Set 1 – DEU | 0.78* | +6.60 | 6.44 | 7.60 | 0.17 |
| Test Set 2 – CR/RANN | 0.87* | +6.39 | 8.57 | 8.56 | 0.14 |
| Test Set 3 - TILDA | 0.65* | -6.97 | 7.52 | 8.42 | 0.22 |

### Brain regions involved in brain age prediction

The voxel-wise method used here to predict brain age resulted in individual coefficient values for each voxel. In order to assess the relative predictive weight for each voxel and to extract the AAL region of interest in which each voxel was situated, the absolute value of the coefficient values were ranked from largest to smallest (using a custom MATLAB function get_beta_labels.m). As a brief overview, the 20 largest negative and positive coefficient values are shown in tables 4 and 5, respectively (see attached excel sheet for coefficient values for all voxels). Voxels with positive coefficient values contributed to older brain age predictions and voxels with negative coefficient values contributed to younger brain age predictions. Figures 1 and 2 show all voxels with negative and positive coefficient values, respectively.

**Table 4.** Overall ranking of coefficient value, coefficient values, MNI coordinates and ROI within the AAL atlas for the 20 largest negative coefficient values.

| Overall Rank | Coefficient values | MNI Coordinates | AAL ROI |
| --- | --- | --- | --- |
| 1 | -0.6085 | [4, -10, 2] | Thalamus_R (aal) |
| 2 | -0.5212 | [42, -10, -6] | Insula_R (aal) |
| 3 | -0.4807 | [38, -46, -42] | Cerebellum_7b_R (aal) |
| 4 | -0.4129 | [18, -34, 0] | Hippocampus_R (aal) |
| 5 | -0.4059 | [18, -34, 2] | Hippocampus_R (aal) |
| 9 | -0.2724 | [-32, -54, -58] | Cerebellum_8_L (aal) |
| 11 | -0.2467 | [-14, 2, 20] | Caudate_L (aal) |
| 12 | -0.2050 | [36, 22, 4] | Insula_R (aal) |
| 17 | -0.1708 | [52, -6, 30] | Postcentral_R (aal) |
| 18 | -0.1692 | [16, -34, 2] | Thalamus_R (aal) |
| 19 | -0.1650 | [44, -4, -12] | Temporal_Sup_R (aal) |
| 21 | -0.1473 | [16, -32, 4] | Thalamus_R (aal) |
| 22 | -0.1427 | [-16, -32, 4] | Thalamus_L (aal) |
| 24 | -0.1374 | [-20, -48, -60] | Cerebellum_8_L |
| 25 | -0.1371 | [8, 12, 42] | Cingulum_Mid_R (aal) |
| 26 | -0.1370 | [-36, -80, -38] | Cerebellum_Crus2_L (aal) |
| 27 | -0.1349 | [8, 34, 26] | Cingulum_Ant_R (aal) |
| 28 | -0.1313 | [-32, -16, -4] | Putamen_L (aal) |
| 29 | -0.1274 | [-6, -16, 44] | Cingulum_Mid_L (aal) |
| 33 | -0.1236 | [18, -42, -50] | Cerebellum_9_R (aal) |

**Table 5.**
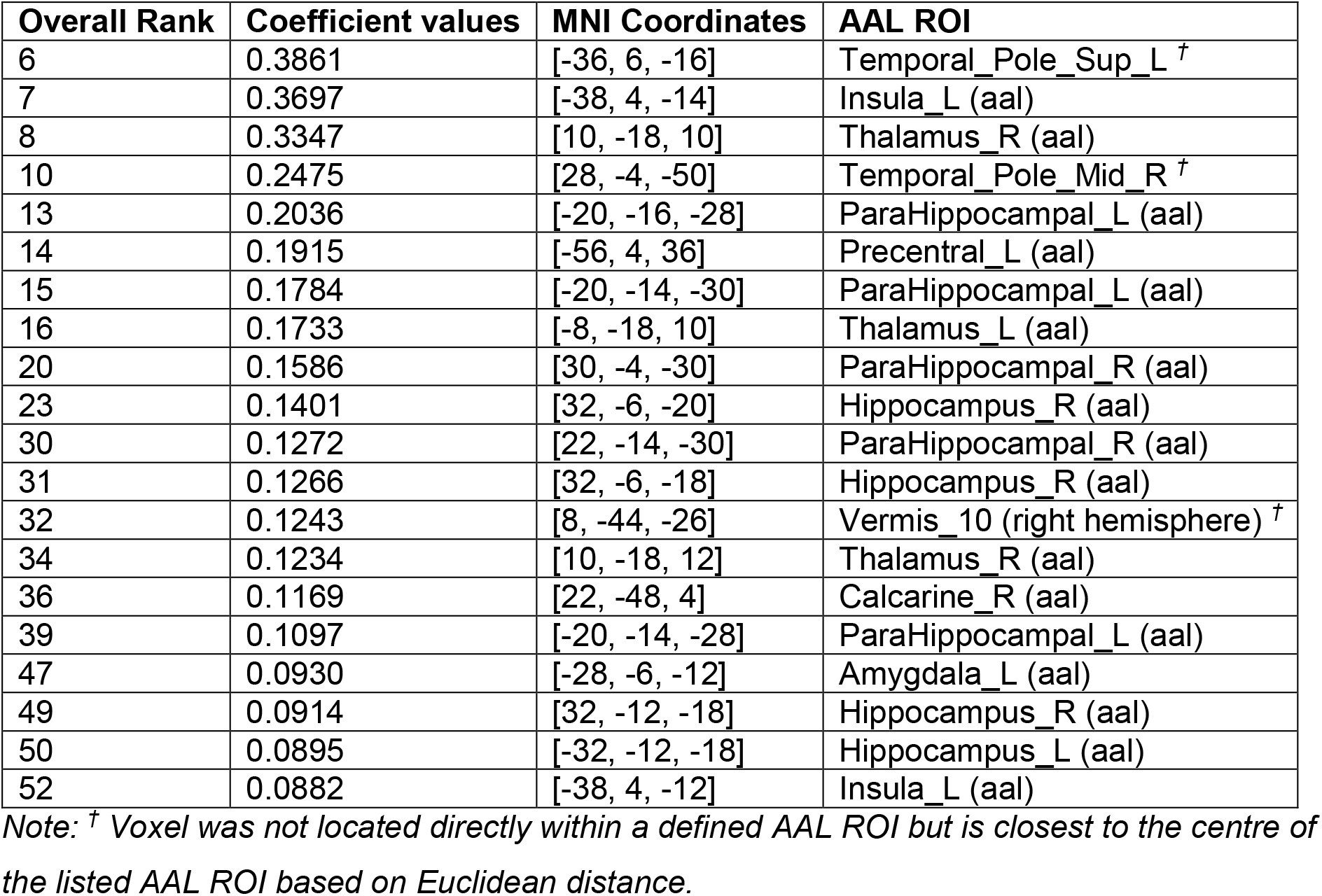
Overall ranking of coefficient value, coefficient values, MNI coordinates and ROI within the AAL atlas for the 20 largest positive coefficient values.

| Overall Rank | Coefficient values | MNI Coordinates | AAL ROI |
| --- | --- | --- | --- |
| 6 | 0.3861 | [-36, 6, -16] | Temporal_Pole_Sup_L <sup>†</sup> |
| 7 | 0.3697 | [-38, 4, -14] | Insula_L (aal) |
| 8 | 0.3347 | [10, -18, 10] | Thalamus_R (aal) |
| 10 | 0.2475 | [28, -4, -50] | Temporal_Pole_Mid_R <sup>†</sup> |
| 13 | 0.2036 | [-20, -16, -28] | ParaHippocampal_L (aal) |
| 14 | 0.1915 | [-56, 4, 36] | Precentral_L (aal) |
| 15 | 0.1784 | [-20, -14, -30] | ParaHippocampal_L (aal) |
| 16 | 0.1733 | [-8, -18, 10] | Thalamus_L (aal) |
| 20 | 0.1586 | [30, -4, -30] | ParaHippocampal_R (aal) |
| 23 | 0.1401 | [32, -6, -20] | Hippocampus_R (aal) |
| 30 | 0.1272 | [22, -14, -30] | ParaHippocampal_R (aal) |
| 31 | 0.1266 | [32, -6, -18] | Hippocampus_R (aal) |
| 32 | 0.1243 | [8, -44, -26] | Vermis_10 (right hemisphere) <sup>†</sup> |
| 34 | 0.1234 | [10, -18, 12] | Thalamus_R (aal) |
| 36 | 0.1169 | [22, -48, 4] | Calcarine_R (aal) |
| 39 | 0.1097 | [-20, -14, -28] | ParaHippocampal_L (aal) |
| 47 | 0.0930 | [-28, -6, -12] | Amygdala_L (aal) |
| 49 | 0.0914 | [32, -12, -18] | Hippocampus_R (aal) |
| 50 | 0.0895 | [-32, -12, -18] | Hippocampus_L (aal) |
| 52 | 0.0882 | [-38, 4, -12] | Insula_L (aal) |

Note: <sup>†</sup> Voxel was not located directly within a defined AAL ROI but is closest to the centre of the listed AAL ROI based on Euclidean distance.

**Fig 1:**
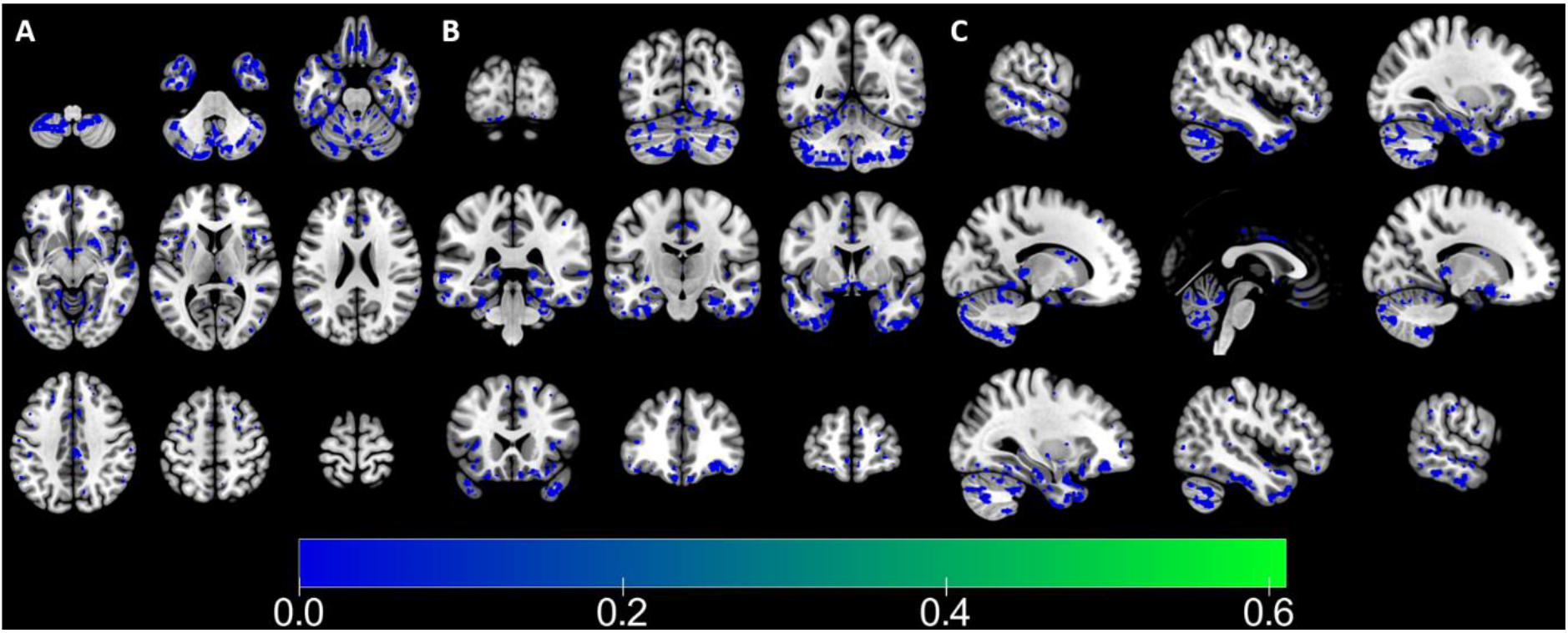
Voxels with negative coefficient values. A = Axial Plane, B = Coronal Plane, C = Sagittal Plane. Note: Signs of these voxel values were flipped to aid plotting.

**Fig 2:**
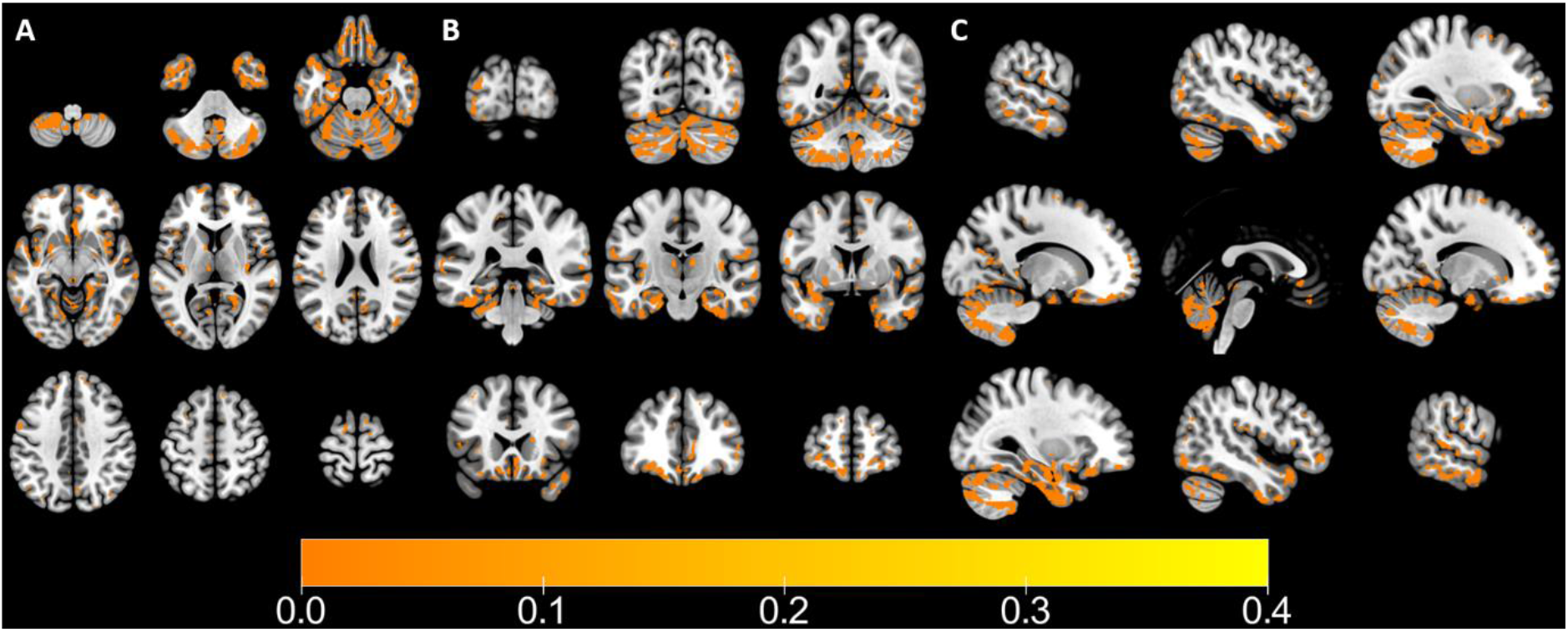
Voxels with positive coefficient values. A = Axial Plane, B = Coronal Plane, C = Sagittal Plane.

### BrainPAD and Cognitive Function

Across multiple datasets, higher brainPAD scores were significantly correlated with reduced performance on measures of general cognitive status, semantic verbal fluency, processing speed, cognitive flexibility, and visual attention (see Figure 3 and Table 6).

**Fig 3:**
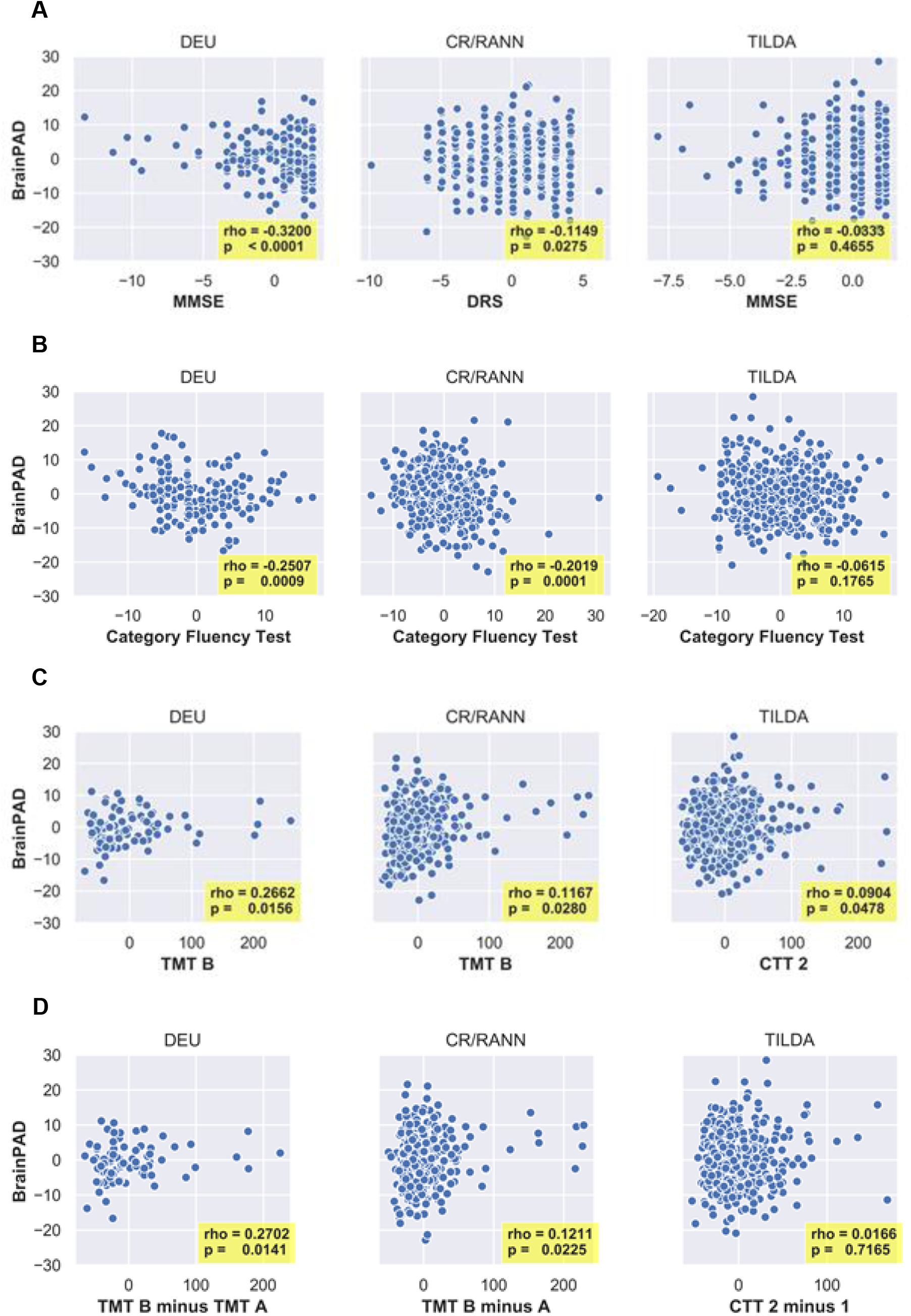
Scatterplots of replicated correlations between the residuals of brainPAD and cognitive measures after regressing brainPAD on age and gender, and each cognitive measure on brainPAD on age and gender. A: General cognitive status; B: Semantic verbal fluency; C: Processing speed, visual attention, and cognitive flexibility, D: Visual attention and cognitive flexibility. For scatterplots of non-replicated correlations, see Supplementary Info, figure S.4.

**Table 6.** Results of Spearman’s partial correlations between brainPAD and 17 cognitive domains.

| Cognitive Domain | DEU |  |  | CR/RANN |  |  | TILDA |  |  | Probability of replicating by chance | Sig. by max statistic correction (where finding not replicated) |
| --- | --- | --- | --- | --- | --- | --- | --- | --- | --- | --- | --- |
|  | rho | df | p | rho | df | p | rho | df | p |  |  |
| General Cognitive Status | -0.3199 | 168 | <0.0001 | -0.1449 | 366 | 0.0275 | 0.0333 | 481 | 0.4655 | <0.00001 | n/a |
| Premorbid Intelligence | n/a |  |  | -0.2322 | 358 | <0.0001 | 0.0485 | 482 | 0.2873 | n/a | CR/RANN |
| Phonemic Verbal Fluency | -0.3259 | 134 | 0.0001 | -0.0771 | 356 | 0.1454 | n/a |  |  | n/a | DEU |
| Semantic Verbal Fluency | -0.2507 | 171 | 0.0009 | -0.2019 | 357 | 0.0001 | -0.0615 | 483 | 0.1765 | <0.00001 | n/a |
| Processing Speed | 0.1232 | 89 | 0.2448 | 0.0595 | 357 | 0.2610 | 0.1208 | 483 | 0.0077 | n/a | None |
| Processing Speed, Visual Attention, Cognitive Flexibility | 0.2662 | 80 | 0.0156 | 0.1167 | 353 | 0.0279 | 0.0904 | 478 | 0.0478 | 0.00005 | n/a |
| Visual Attention, Cognitive Flexibility | 0.2702 | 80 | 0.0141 | 0.1211 | 353 | 0.0225 | 0.0166 | 478 | 0.7165 | 0.00097 | n/a |
| Cognitive Flexibility | 0.0722 | 46 | 0.6258 | 0.0429 | 323 | 0.4411 | n/a |  |  | n/a | None |
| Response Inhibition, Selective Attention | 0.0854 | 146 | 0.3019 | -0.1755 | 355 | 0.0009 | n/a |  |  | n/a | None |
| Sustained Attention (Errors of Commission) | n/a |  |  | 0.0203 | 172 | 0.7902 | 0.0499 | 478 | 0.2752 | n/a | None |
| Sustained Attention (Reaction Time) | n/a |  |  | -0.0212 | 172 | 0.7813 | 0.0436 | 475 | 0.3425 | n/a | None |
| Verbal Episodic Memory (Immediate) | 0.2194 | 171 | 0.0037 | -0.0407 | 356 | 0.4428 | -0.0347 | 483 | 0.4114 | n/a | None |
| Verbal Episodic Memory (Delayed) | 0.2797 | 171 | 0.0002 | 0.0343 | 356 | 0.5173 | 0.0122 | 483 | 0.7887 | n/a | None |
| Verbal Episodic Memory (Learning) | -0.3196 | 171 | <0.0001 | 0.0657 | 356 | 0.2151 | n/a |  |  | n/a | DEU |
| Working Memory | -0.1310<br>-0.2974 | 167<br>166 | 0.0895<sup>†</sup><br>0.0001<sup>‡</sup> | -0.0469 | 360 | 0.3759 | n/a |  |  | n/a | None |
| Visuospatial Ability | -0.0809 | 76 | 0.4815 | -0.1824 | 352 | 0.0006 | n/a |  |  | n/a | CR/RANN |

Note: <sup>†</sup> Digit Span Forwards; <sup>‡</sup> Digit Span Backwards

## Discussion

A penalized regression approach was able to produce accurate brain-age predictions from T1 MRI data in three independent datasets. In non-demented adults, brain predicted-age difference (brainPAD), calculated by subtracting these brain-age predictions from chronological age, was negatively correlated with general cognitive status, semantic verbal fluency, processing speed, visual attention, and cognitive flexibility; and visual attention and cognitive flexibility in multiple datasets. BrainPAD was significantly correlated with phonemic verbal fluency, premorbid intelligence, verbal episodic memory (learning score), and visuospatial ability in single datasets after controlling for multiple comparisons; however, these correlations were not replicated in another dataset so we do not have strong evidence here in support of these relationships. BrainPAD was not significantly correlated with processing speed, cognitive flexibility, response inhibition and selective attention, sustained attention, verbal episodic memory (immediate recall or delayed recall), or working memory in any dataset.

### BrainPAD and Cognition

#### General Cognitive Status

BrainPAD was negatively correlated with general cognitive status, as measured using the MMSE and DRS, in DEU and CR/RANN, and the replication of this result across both datasets was statistically significant. However, brainPAD was not significantly correlated with the MMSE in TILDA. Nonetheless, given the statistically significant replication across two of the three datasets, there is reliable evidence in support of the correlation between brainPAD and general cognitive status in healthy older adults. Previous studies have reported that brainPAD is related to general cognitive status, albeit in samples including individuals with MCI, AD, or dementia (Beheshti et al., 2018; Kaufmann et al., 2018), and without adjusting for the effect of age or controlling for multiple comparisons (Beheshti et al., 2018; Cole, Underwood, et al., 2017; but see Table 1 Footnote 4 for information about adjusting for age in Cole, Underwood et al., 2017). In contrast to our findings, Gaser *et al*. (2013) reported that brainPAD was correlated with the CDR and ADAS but not the MMSE in an MCI sample. However, Gaser *et al*. (2013) did not account for the effect of age. While Löwe *et al*. (2016) reported that brainPAD was negatively correlated with the MMSE across mixed samples of APOE e4 carriers and non-carriers (including healthy controls, MCI, and AD), it was not significantly correlated with the MMSE within healthy control and MCI subgroups. Sample sizes within these subgroups were relatively small, ranging from 14 to 81 participants. Consequently, the correlations between brainPAD and the MMSE in these participants may not have been adequately powered to reach significance. Our study is the first to report a relationship between brainPAD and measures of general cognitive status in healthy adults while controlling for the effects of age and correcting for multiple comparisons. This is also the first study to investigate the relationship between brainPAD and DRS score. Our findings provide strong support for the existence of a significant negative relationship between brainPAD and general cognitive status.

It was beyond the scope of the present study to formally test associations between brain areas contributing to brainPAD and cognitive measures. The voxels that best predicted chronological age were found in the thalamus, hippocampus, parahippocampal gyrus and amygdala. There is previous evidence of positive correlations between the MMSE and GM volume within the amygdala, hippocampus, parahippocampal gyrus (Dinomais et al., 2016), and the thalamus (Ferrarini et al., 2008; Yi et al., 2016), and between the DRS and combined GM and WM volume within the left hippocampus (Fama et al., 1997). In our brainPAD model, voxels within these areas were among the 20 voxels most heavily contributing to brainPAD. However, it is more appropriate to consider brainPAD as a summary score representing global brain atrophy as it was driven by contributions from voxels spread across the brain. Impaired performance on the MMSE is also associated with global brain (Fox, Scahill, Crum, & Rossor, 1999; Mak et al., 2015) and GM atrophy (Brunetti et al., 2000). Similarly, there is a positive relationship between DRS and total cortical GM volume (Stout, Jernigan, Archibald, & Salmon, 1996). As such, the correlation between brainPAD and general cognitive status is supported by previous evidence of brain changes related to performance on these measures. This strengthens the argument in favour of brainPAD as an objective measure of general cognitive function given that brainPAD is not subject to the various biases and effects (e.g. low reliability, practice effects) that limit the MMSE (Galasko, Abramson, Corey-Bloom, & Thal, 1993; Pfeffer, Kurosaki, Chance, Filos, & Bates, 1984; Tombaugh & McIntyre, 1992) and the DRS (Emery, Gillie, & Smith, 1996; Green, Woodard, & Green, 1995).

#### Semantic Verbal Fluency

BrainPAD was significantly negatively correlated with semantic verbal fluency, as measured using the Animals task, in both DEU and CR/RANN but not in TILDA. Regardless, the replication of this result across both DEU and CR/RANN was statistically significant. This finding contradicts non-significant correlations between brainPAD and composite measures of semantic and phonemic verbal fluency (Cole, Underwood, et al., 2017; Richard et al., 2018), although the former study used age-adjusted t-scores to control for the age-cognition relationship rather than adding age as a covariate to the brainPAD-fluency measure (cf. Le et al., 2018). As semantic verbal fluency is associated with age (Clark et al., 2009; Santos Nogueira, Azevedo Reis, & Vieira, 2016), the failure to adjust for age may have obscured a significant effect. Alternatively, these previously reported non-significant correlations could be explained by the use of composite measures of both phonemic and semantic fluency as we did not find strong evidence for a relationship between phonemic verbal fluency and brainPAD (although it was significant in DEU, this correlation was not replicated in CR/RANN). Therefore, it is possible that a non-significant relationship between phonemic fluency and brainPAD in the Cole *et al*. (2017) and Richard *et al*. (2018) study may have diluted a possible significant relationship between semantic fluency and brainPAD. In a study controlling for age, brainPAD was found to significantly negatively correlate with semantic verbal fluency (Franke et al., 2013). The brainPAD-semantic verbal fluency correlation is further supported by evidence that some voxels that contribute heavily to brainPAD in our model are located in regions where GM volume has been positively correlated with semantic verbal fluency, including the cerebellum (Grogan, Green, Ali, Crinion, & Price, 2009) and the thalamus, in adolescents born prematurely (Giménez et al., 2006), and in regions where lesions have been negatively related to semantic verbal fluency, such as the left temporal lobe and insula (Baldo, Schwartz, Wilkins, & Dronkers, 2006). Although the Animals task has been described as an optimal test of neuropsychological function (Ardila, Ostrosky-Solís, & Bernal, 2006), scores on this task are affected by various factors, including scoring and administration procedures (Woods, Wyma, Herron, & Yund, 2016) and practice effects (Cooper et al., 2001; Harrison, Buxton, Husain, & Wise, 2000; Wilson, Watson, Baddeley, Emslie, & Evans, 2000). As such, brainPAD, as an objective marker of general brain health and global cognitive function, could be a viable alternative to the Animals task. In sum, our results provide further evidence in support of a correlation between brainPAD and semantic verbal fluency.

#### Processing speed, visual attention, and cognitive flexibility

Across all three datasets, brainPAD was negatively correlated with processing speed, visual attention, and cognitive flexibility as measured by trail-making tests (TMT B or CTT 2). The TMT B is a relatively sensitive measure of cognitive decline: completion times were shown to be significantly different between healthy controls, MCI, and AD (Ashendorf et al., 2008). Likewise, the CTT 2 is sensitive to cognitive decline, with differences between AD and healthy controls (Lin et al., 2014), and between healthy controls, MCI, and AD (Guo et al., 2010). Therefore, it is no surprise that processing speed, visual attention, and cognitive flexibility were also negatively correlated with an index of accelerated brain ageing. Indeed, previous studies have reported similar results for trail-making versus brainPAD; however, these studies did not correct for multiple comparisons (Cole, Underwood, et al., 2017) or used clinical samples (TBI; Cole et al., 2015). Our data therefore augment these findings by replicating this result across three independent datasets. Tentative neurobiological support can be found for this correlation in the overlap in correlations between TMT B performance and GM volume in regions containing voxels which contribute heavily to brainPAD in our model. TMT B performance is negatively correlated with medial temporal lobe atrophy (Oosterman et al., 2010) and is positively related with GM density within the cerebellum in patients with spinocerebellar ataxia (Rentiya, Khan, Ergun, Ying, & Desmond, 2017). This evidence could suggest that brainPAD may be a potential objective measure of cognitive decline as it is not subject to the same factors which bias trail-making performance, including to practice effects (Bartels, Wegrzyn, Wiedl, Ackermann, & Ehrenreich, 2010), rater effects (Feeney et al., 2016) and participant literacy (Vaucher et al., 2014).

#### Visual attention and cognitive flexibility

BrainPAD was also negatively correlated with visual attention and cognitive flexibility (TMT B minus A), in DEU and CR/RANN, but not in TILDA (CTT 2 minus 1). Replication of this finding (albeit with relatively small rho values) in DEU and CR/RANN suggests a modest association between visual attention and cognitive flexibility. The relationship between brainPAD and TMT B minus A was only investigated in one previous study, in a TBI sample, (Cole et al., 2015) where a significant positive correlation was reported. Task-based fMRI studies have shown significantly increased activation during TMT B versus TMT A within the left precentral gyrus (Moll, Oliveira-Souza, Moll, Bramati, & Andreiuolo, 2002) and the insula (Zakzanis, Mraz, & Graham, 2005). Additionally, a voxel-based lesion-symptom mapping study reported a negative association between lesions within the left insular cortex and a TMT task accuracy score, which like the TMT B minus A score attenuates the influence of processing speed and reflects cognitive flexibility (Varjačić et al., 2018). Like the TMT B score, the TMT B minus A is also correlated with atrophy of the bilateral medial temporal lobes (Oosterman et al., 2010). All of these regions contain voxels which contribute to older brain age in our model and therefore these various findings may provide some neurobiological basis for the correlation between brainPAD and visual attention and cognitive flexibility. As such, although the TMT B minus A can distinguish between stable and progressive MCI on a group level (Zanetti et al., 2006), and is associated with reduced mobility, increased mortality risk (Vazzana et al., 2010) and slower walking speed (Ble et al., 2005), as a derived measure of the TMT, the TMT B minus A index is similarly affected by the various factors that can limit interpretation of the TMT B scores. Therefore, given the correlation shown here between TMT B minus A and brainPAD, brainPAD may be a potential objective measure of general cognitive function.

It is notable that several significant brainPAD-cognition relationships were observed in the DEU and CR/RANN datasets, but not in TILDA. We tentatively offer some suggestions for this pattern of results. Confounding factors obscuring the brainPAD-general cognitive status relationship may have been uniquely present in TILDA. Whereas the DEU and CR/RANN cohorts were part of neuroimaging research studies, which have typically strict inclusion criteria, the TILDA MRI sample were a subset of a large nationally representative longitudinal study encompassing health, economic and social research (B. J. Whelan & Savva, 2013). TILDA therefore had few inclusion criteria: being at least 50 years old, having a residential address, and absence of dementia at baseline (Kearney et al., 2011; Savva, Maty, Setti, & Feeney, 2013). TILDA’s MRI sample were screened for MRI contraindications and were on average healthier than the full sample, but it is likely that the TILDA sample included participants who might normally be excluded from neuroimaging research studies (e.g., those using psychotropic or cardiovascular medication). Moreover, the range of some cognitive measures in TILDA was also smaller than DEU and CR/RANN in some cases (see Supplemental Table 3): notably for general cognitive status, and visual attention and cognitive flexibility, where the brainPAD-cognition correlations were not replicated within TILDA. Restricted range of scores on these measures in TILDA may have contributed to smaller correlation coefficients (Bland & Altman, 2011; Mendoza & Mumford, 1987). Additionally, the age range within TILDA was smaller than both DEU and CR/RANN which may have reduced the statistical power of the brainPAD-cognition correlations within TILDA as range restriction on covariates has also been shown to reduce power (Miciak, Taylor, Stuebing, Fletcher, & Vaughn, 2016) and decrease the magnitude of correlation coefficients (Sackett & Yang, 2000).

### Model evaluation

We evaluated our model based on its predictive accuracy in three independent test sets, as proposed by Madan and Kensinger (2018). While internal cross-validation is a valuable and widely used technique that can attenuate overfitting (Arlot & Celisse, 2010); the use of cross-validation in certain situations and when it is not implemented correctly, can result in overestimated prediction accuracy and overfitting (Saeb, Lonini, Jayaraman, Mohr, & Kording, 2016; Skocik, Collins, Callahan-Flintoft, Bowman, & Wyble, 2016; Varoquaux et al., 2017). For brainPAD to be considered for clinical use, it must perform accurately with MRIs acquired in different scanners and under different protocols. However, in most instances of cross-validation, while the test set is split and held completely independent from the training set, factors common to both sets, such as scanner and protocol, could influence model performance. As such, the gold-standard evaluation for brainPAD should be accurate performance on independent external datasets.

The significant correlations between chronological age and brain-predicted age in all three external datasets shows that our model is accurate and generalizable (0.65, 0.78, and 0.87 for external datasets). Although the magnitude of these correlations is lower than correlations reported elsewhere, ranging from 0.91 to 0.94 (Cole et al., 2015; Cole, Poudel, et al., 2017; Franke et al., 2010; Lancaster et al., 2018; Liem et al., 2017), it exceeds other externally validated brain-predicted age studies, ranging from 0.65 to 0.85 (Beheshti et al., 2018; Madan & Kensinger, 2018; Varikuti et al., 2018).

With respect to mean absolute error (MAE), our model did not perform as well as other externally validated studies, ranging from 4.28 to 7.5 years (Beheshti et al., 2018; Cole, Ritchie, et al., 2018; Franke et al., 2010; Lancaster et al., 2018; Madan & Kensinger, 2018). As we targeted an interpretable model of brain age, we may have lost some precision by not integrating WM information as input in the model, as was done by Cole *et al*. and Franke *et al*. Another potential reason is that other studies centered the age predictions using the mean of the ages from the test set. Although this correction is typically not explicitly described in method sections, Madan and Kensinger (2018) note that this is a standard correction in brain age prediction. Moreover, some studies also match the variance in predicted age in the test set with the variance of the training data (Madan & Kensinger, 2018). Both corrections are principled and acceptable methods of correcting for the regression to the mean artefact in brain age predictions but they result in biased age predictions in the test set. These corrections also limit the use of brainPAD to make single subject predictions, as both the test set mean and variance are used in the prediction. Our method used only training set information and therefore produced slightly less accurate but less biased predictions. Finally, our model may also appear to be less precise in terms of MAE as an artefact of the greater age range of our sample in comparison to most brainPAD studies. An alternative metric, the weighted MAE (calculated by dividing the MAE by the age range of the sample), may enable better comparisons across studies with different age ranges (Cole, Franke, et al., 2018). While our weighted MAE is higher than some studies, ranging from 0.072 to 0.087 (Lancaster et al., 2018; Liem et al., 2017), the lowest weighted MAE in our sample (0.14 in CR/RANN) outperformed this metric when calculated for other studies, 0.178 (Beheshti et al., 2018), and 0.18 (Varikuti et al., 2018) and is comparable to 0.139 (Franke et al., 2010, ‘Test 4’ external test set). As such, the predictive accuracy of our model is comparable to the rest of the literature and is arguably less biased as only training set information is used.

### Model interpretation

#### Model interpretability

The interpretability of machine learning models is an important and widely discussed problem (Doshi-Velez & Kim, 2017), and although it is poorly defined (Lipton, 2018) it has been described as *“the ability to explain or to present in understandable terms to a human”* (Doshi-Velez & Kim, 2017, p. 2) and elsewhere as the ability to “*understand the contribution of individual features in the model*” (Lou, Caruana, & Gehrke, 2012, p. 1). Additionally, Lipton (2018) argued that for a model to be considered truly interpretable, it should possess the following three properties: *algorithmic transparency* (i.e. it should be possible to understand the mechanism by which the model works), *decomposability* (each part of the model, such as the model input and parameters, should have an intuitive explanation), and *simulatability* (a person should be able to consider the entire model at once). We contend that our model possesses these three properties as well as conforming to the definitions proposed above. First, our model possesses algorithmic transparency in that the Elastic Net is a penalized linear regression. Second, our model possesses decomposability. The inputs to the model were GM voxel density values and the parameters, or beta coefficient values, weighted the contribution of each individual value to the model output, which is brain predicted age. Third, our model possesses simulatability as the entire model can be considered as follows: summing the multiplication of GM voxel density values by the average contribution of these voxels to the prediction of chronological age in the training set (i.e., the beta coefficient values) resulted in a prediction of a new individual’s brain age.

#### Brain regions involved in brain age prediction

In addition to the good interpretability of our model, the model is also biologically plausible as the top 20 brain voxels contributing to older brain age are all located in brain regions previously shown to be vulnerable to age-related GM volumetric decline, including the insula (Good et al., 2001; Kennedy et al., 2009; Resnick, Pham, Kraut, Zonderman, & Davatzikos, 2003; Taki et al., 2011), thalamus (Abe et al., 2008; Sullivan, Rosenbloom, Serventi, & Pfefferbaum, 2004; Taki et al., 2011; Walhovd et al., 2005), and temporal cortex (Abe et al., 2008). Within the temporal cortex specifically, there were strong contributions to older brain age from voxels in the temporal pole, where there are negative ageing effects on GM volume (Allen, Bruss, Brown, & Damasio, 2005; Lemaitre et al., 2012). Voxels within temporal lobe structures, such as the amygdala, hippocampus and parahippocampal gyrus also strongly contributed to older brain age which is in line with evidence of age-related volumetric declines in the left amygdala (Giorgio et al., 2010), bilateral hippocampus (Giorgio et al., 2010; Jernigan et al., 2001) and bilateral parahippocampal gyrus (Jernigan et al., 2001; Taki et al., 2011). Similarly, another heavily weighted contribution to brain age was found from a voxel within the left precentral gyrus, where GM volume is negatively associated with age (Kennedy et al., 2009; Taki et al., 2011). Finally, there was a strong contribution from a voxel within the cerebellar vermis, where there are increased rates of GM volume loss during ageing in comparison to other areas of the cerebellum (Yu, Korgaonkar, & Grieve, 2017).

The voxels which contributed most strongly to an older brain age were located within regions vulnerable to age-related GM volumetric decline. However, some of the voxels that contributed most strongly to a younger brain age were found adjacent to the negatively weighted voxels. One potential explanation is that brainPAD measures GM atrophy, and therefore individuals with relatively less atrophy will have higher GM density on the edges of the cortex and subcortical structures. In contrast, individuals with more atrophy will have greater GM density distal from the edges of the cortex (when registered in standard space), and consequently positively weighted voxels (i.e., which contributed to older brain age) were located in those distal regions.

### Limitations

A possible limitation of the current model is that it uses only voxel-wise GM density data and thus our model may have lower accuracy due to this restricted feature set. Other brain age models have used feature sets including combinations of cortical and subcortical GM regional volumes (Steffener et al., 2016); combinations of GM voxel density values, cortical thickness, and regional volume data (Gutierrez Becker et al., 2018); combinations of cortical thickness, cortical surface area, subcortical volume, and functional connectivity information (Liem et al., 2017); and combinations of GM and WM voxel-wise density information (Cole et al., 2015; Cole, Ritchie, et al., 2018; Cole, Underwood, et al., 2017). However, such feature sets typically require dimension reduction such as PCA (Gutierrez Becker et al., 2018) or even dot products to combine GM and WM data (Cole et al., 2015; Cole, Ritchie, et al., 2018; Cole, Underwood, et al., 2017). These steps reduce the interpretability of the relationship between the original feature and brain age. Our aim was to produce an interpretable model, an aim which required a simple feature set. While this focus on improved interpretability may have limited our model’s accuracy as larger and more complex feature sets often produce more accurate predictions (Scheinost et al., 2019), our model’s accuracy is still comparable to other models reported to-date in the literature. Likewise, while our model is readily interpretable; greater interpretability could potentially be achieved by forcing sparsity to limit the number of voxels making significant contributions to brain age predictions. Modified Elastic Net algorithms, such as Enet-BETA (Liu & Li, 2017), can obtain sparser models which would reduce the number of predictive voxels, thereby further improving interpretability. However, as the Elastic Net’s prediction accuracy can increase with feature set size (Jollans et al., in revision), limiting the feature set size could reduce model accuracy. Our model may strike the right balance between interpretability and accuracy.

The major limitation of our study is that for the majority of the cognitive domains investigated here, we used different cognitive measures to assess the putatively same cognitive processes. For example, although we considered the CTT 2 as a direct ‘culture-free’ analogue of the TMT B, as it is widely described (Elkin-Frankston, Lebowitz, Kapust, Hollis, & O’Connor, 2007; Messinis, Malegiannaki, Christodoulou, Panagiotopoulos, & Papathanasopoulos, 2011), the CTT 2 has different stimuli (shapes and colors vs numbers and letters) and takes longer because it has more stimuli (Mitrushina, Boone, Razani, & D’Elia, 2005). Consequently, some have argued, based on findings of significant difference in mean scores on CTT 2 and TMT B, that the tests are not direct equivalents (Dugbartey, Townes, & Mahurin, 2000; Strauss, Sherman, & Spreen, 2006). However, mean scores for both measures are calculated as time to completion and thus a difference in means between both measures reflects a difference primarily in test length. A more appropriate measure of test equivalence would be correlations between mean scores, and various studies report significant correlations between both measures (Dugbartey et al., 2000; Elkin-Frankston et al., 2007; Lee, Cheung, Chan, & Chan, 2000; Messinis et al., 2011). Similar arguments might be made for the other tests (e.g. the MMSE and DRS) that we used to assess the same cognitive constructs (e.g. general cognitive status). While it would be preferable to use the identical measures across datasets, our study used existing data and was designed after data collection. As a result, this approach was not possible here. Nonetheless, the measures used here were broadly comparable in that they are apparent measures of the same underlying cognitive constructs and it is these constructs which we are most interested in, more so than the actual measures.

## Conclusion

The brain age model presented here is accurate and generalizable as it significantly predicts chronological age in 3 independent datasets. Furthermore, this model is interpretable and biologically plausible as older brain age is driven by decreased GM density in voxels that have been previously shown to be vulnerable to GM atrophy and volume loss. Finally, brainPAD scores, calculated using this model, are associated with reduced cognitive performance within the domains of general cognitive status; semantic verbal fluency; processing speed, visual attention, and cognitive flexibility; and visual attention and cognitive flexibility. The replication of these correlations in multiple datasets demonstrates that the relationship between brainPAD and these domains of cognitive function is robust to cultural- and site/scanner effects. As such, given that brainPAD is also not limited by task effects which can hinder neuropsychological assessment, these findings provide support for the use of brainPAD as an objective measure of general cognitive function with applications as a general measure of brain health and cognitive performance in the clinic and as a summary outcome measure for intervention studies in research settings.

## Funding Acknowledgements

RB is supported by the Irish Research Council under grant number EPSPG/2017/277. LRD and RW are supported by the Science Foundation Ireland under grant number 16/ERCD/3797. RR is supported by a PhD scholarship funded by the Region Calabria. Data collection in Dokuz Eylul University, managed and supervised by GGY and DDS, was partially supported by the Turkish National Science and Research Council (TUBITAK, Grant number: 112S459) and the Dokuz Eylul University Scientific Research Projects (Grant number: 2018.KB.SAG.084). The Irish Longitudinal Study on Ageing is funded by core grants from the Health Research Board, Atlantic Philanthropies and Irish Life. MRI data collection in TILDA was supported by the Centre for Advanced Medical Imaging (CAMI) at St. James’ Hospital, Dublin. IHR thanks The Atlantic Philanthropies for their grant to the Global Brain Health Institute. YS is supported by NIA RF1 AG038465 and R01 AG026158. The funding agencies had no involvement in the conduct of the research or preparation of the article.

