## Supplemental Information for "Brain-predicted age difference score is related to specific cognitive functions: A multi-site replication analysis"

### Supplementary Methods

#### MRI pre-preprocessing

All scans were automatically reoriented using a custom MATLAB function, auto_reorient.m - based on the same-named function created by Carlton Chu, which aligned each individual image with the MNI single subject T1 scan. In the training set, badly oriented scans, or scans with gross artefacts, were excluded from further analysis. Although this reduced the training set sample size, this eliminated the need for time-consuming manual re-orientation of individual images. To prevent data loss in the test set, any badly oriented scans were manually re-oriented but scans with artefacts were still excluded.

#### Machine Learning

The Elastic Net combines the Least Absolute Shrinkage and Selection Operator (LASSO) regression, where regression weights are penalised for increasing model complexity based on their absolute size and can be set to zero, and ridge regression, where regression weights are penalised for increasing model complexity based on their squared values and as such cannot be set to zero (Zou & Hastie, 2005). The Elastic Net is particularly well-suited for data with a much larger number of predictors than observations, such as neuroimaging data.

Each nested training set was divided into 10 CV folds, each consisting of 10% of the subsampled training set. Nine folds were then used to create the regression model and the model's prediction were then tested on the one left-out fold. This entire procedure was repeated 10 times, with each CV fold being left-out once. Furthermore, within each fold, nested cross-validation with 10 partitions was then used for optimisation of the two Elastic Net model parameters: alpha ($\alpha$), which is the weight of the lasso vs. ridge regularization and lambda ($\lambda$), which is the regularization coefficient. Thirty values of each parameter were used, with $\alpha$ parameters ranging from 1e-25 to $\lambda$ ranging from 1 to 1e-04. The most frequently occurring parameter values across nested CV folds were used to create the final prediction model for each CV fold.

The data resampling ensemble approach used in this model, which created 25 nested training sets with a 50:50 gender ratio, controlled for the effect of gender and reduced any possible individual model effects.

***Table S.1***

*List of open-access repositories used with information on exclusion during image processing, age and gender of dataset, and scanner information.*

| **Dataset** *(Country - site)* | **Eligible n** | **Excluded n** | **Reason for exclusion** | **Final n** | **Mean age of final n** (SD, range) | **Male/ Female** | **Scanner** (field strength) | **Voxel dimensions** (mm) |
|---|---|---|---|---|---|---|---|---|
| **Autism Brain Imaging Data Exchange (ABIDE)** | 102 | 39 | Failed auto reorientation | 63 | 28.70 (11.32, 18 - 64) | 46/17 | | |
| ***USA – Barrow Neurological Institute*** | | | | | | | Philips Ingenia (3T) | 1.11 x 1.11 x 1.2 |
| ***USA – Indiana University*** | | | | | | | Siemens TriTim (3T) | 0.7 x 0.7 x 0.7 |
| ***France – Institut Pasteur and Robert Debré Hospital*** | | | | | | | Phillips Achieva (1.5T) | 1.00 x 1.00 x 1.00 |
| ***Belgium – Katholieke Universiteit Leuven*** | | | | | | | Phillips Achieva Ds (3T) | 1.20 x 1.20 x 1.20 |
| ***SA – New York University Langone Medical Center*** | | | | | | | Siemens Allegra (3T) | 1.30 x 1.00 x 1.30 |
| ***Ireland – Trinity College Dublin*** | | | | | | | Philips Intera Achieva (3T) | 0.89 x 0.89 x 0.89 |

| | | | | | | | | |
|---|---|---|---|---|---|---|---|---|
| **The Neuro Bureau – Berlin: Mind & Brain *(Germany)*** | 49 | 22 | Failed auto reorientation | 27 | 32.62 (6.18, 22.24 – 49.37) | 15/12 | Siemens TrioTim (3T) | 1 x 1 x 1 |
| **Beijing Normal University *(China)*** | 179 | 10 | Failed auto reorientation | 169 | 21.22 (1.88, 18 – 28) | 66/103 | Siemens TrioTim (3T) | 1.33 x 1.0 x 1.0 |
| **Cleveland Clinic Foundation (CCF) *(USA – Cleveland Clinic Hospital)*** | 31 | 31 | Failed auto reorientation (10); Failed QC (21) | 0 | n/a | n/a | Siemens Trio Tim (3T) | 2 x 1 x 1.2 |
| **Center for Biomedical Research Excellence (COBRE) *(USA – The Mind Research Network)*** | 72 | 13 | Failed auto reorientation | 59 | 34.98 (11.61, 18 – 62) | 38/21 | Siemens Trio Tim (3T) | 1 x 1 x 1 |
| **Dallas Lifespan Brain Study (DLBS) *(USA – University of Texas at Dallas)*** | 315 | 138 | Failed auto reorientation | 177 | 46.82 (18.39, 20.57 – 88.36) | 45/132 | Philips (3T) | |
| **Information eXtraction from Images (IXI)**<br><br>***UK – Hammersmith Hospital***<br><br>***UK –Guy's Hospital***<br><br>***UK – Institute of Psychiatry)*** | 565 | 325 | Failed auto reorientation | 240 | 47.20 (16.17, 19.98 – 80.17) | 62/178 | Philips Medical Systems Intera (3T)<br><br>Philips Gyroscan Intera (1.5T)<br><br>General Electric Signa (1.5T) | 0.9375 x 0.9375 x 1.2 |

| | | | | | | | | |
|---|---|---|---|---|---|---|---|---|
| **Nathan Kline Institute -Rockland Sample (NKI)** ***(USA – Nathan Kline Institute)*** | 143 | 28 | Failed auto reorientation (27); Failed QC (1) | 115 | 42.83 (17.95, 18 - 83) | 40/75 | Siemens TrioTim (3T) | 1 x 1 x 1 |
| **Southwest University Adult Lifespan Dataset (SALD) (*China – Southwest University)*** | 494 | 46 | Failed auto reorientation (26); Failed QC (20) | 448 | 44.73 (17.44, 19 - 80) | 162/286 | Siemens Trio Tim (3T) | 1 x 1 x 1 |
| **Power et al (2014)** ***(USA – Washington University in St. Louis)*** | 83 | 22 | Failed auto reorientation | 61 | 24.56 (2.29, 19.69 – 37.73) | 30/31 | Siemens Trio Tim (3T) | 1 x 1 x 1 |
| **Training set total** | 2033 | 674 | Failed auto reorientation (632); Failed QC (42) | 1359 | 40.04 (17.78, 18 – 88.36) | 504/855 | N/A | N/A |

*Note: Eligible N = Healthy controls >= 18 years old with age and gender data available. All datasets, except for the IXI dataset were downloaded from the 1000 Functional Connectomes Project via the NITRC repository http://fcon_1000.projects.nitrc.org/. The IXI dataset was downloaded from http://brain-development.org/ixi-dataset/.*

#### Cognitive function measures

##### *General Cognitive Status*

Total scores on the mini-mental state examination (MMSE; Folstein, Folstein and McHugh, 1975) in DEU and TILDA, and on the Mattis Dementia Rating Scale-2 (DRS; Jurica, Leitten, & Mattis, 2001) in CR/RANN were used to assess general cognitive status. Both scales assess cognitive functioning across domains which are typically affected by Alzheimer's disease and cognitive impairment (Monsch et al., 1995; Tombaugh & McIntyre, 1992).

##### *Premorbid Intelligence*

Raw scores on the American National Adult Reading Test (AMNART; Grober and Sliwinski, 1991) in CR/RANN and on the National Adult Reading Test (NART; Nelson and Willinson, 1982) in TILDA were used to assess premorbid intelligence. In TILDA, 60.57% of those with NART scores completed the full NART whereas 39.22% completed the first half of the NART only. Participants only proceeded to the second half of the NART if they scored over 20 on the first half. This is both a time-saving measure and serves to reduce distress and anxiety in people with poor reading skills (Strauss, Sherman, & Spreen, 2006). Scores of 0-11 were used as full scores but scores of 12-20 were corrected using a conversion table outlined by Beardsall and Brayne (1990). There was no comparable measure of premorbid intelligence in DEU.

##### *Phonemic Verbal Fluency*

Phonemic verbal fluency was assessed using the total score on a Turkish language version of the FAS test in DEU, the KAS test (Tumac, 1997), and on the CFL test in CR/RANN. These tests measure the ability to spontaneously produce words beginning with specific letters (i.e. 'K' in KAS or 'C' in CFL; Strauss, Sherman and Spreen, 2006). There was no comparable measure of phonemic verbal fluency in TILDA.

##### *Semantic Verbal Fluency*

Semantic verbal fluency was assessed using the total score on the Animals test in all three datasets. A Turkish language version of this task was used in DEU (Tumac, 1997). This test measures the ability to spontaneously produce the name of animals (Strauss et al., 2006).

##### *Processing Speed*

Cognitive processing speed was assessed using time to the Trail Making Task A (TMT; Reitan, 1955) in DEU and CR/RANN, and the Colour Trails Task 1 (CTT; D'Elia *et al.*, 1996) in TILDA. The CTT is considered a cross-culturally valid form of the TMT (Strauss et al., 2006).

*Processing Speed, Visual Attention, and Cognitive Flexibility*

Cognitive processing speed, visual attention, and cognitive flexibility were assessed using time to complete the TMT B in DEU and CR/RANN and the CTT 2 in TILDA.

*Visual Attention and Cognitive Flexibility*

A purer measure of cognitive flexibility and visual attention was obtained by subtracting the simpler TMT A and CTT 1 from the more complex TMT B and CTT 2, respectively. This difference score was calculated in all 3 datasets and controls for general processing speed (Strauss et al., 2006).

*Cognitive Flexibility*

Cognitive flexibility was also measured by the percentage of perseverative errors on the Wisconsin Card Sorting Task (WCST; Heaton, Chelune, Talley, Kay, & Curtiss, 1993) in DEU and CR/RANN.

*Response Inhibition and Selective Attention*

The Stroop test in DEU and CR/RANN was used to assess response inhibition and selective attention. A Turkish version of the Stroop test, the Stroop Test Çapa Version, was used in DEU (Emek-Savaş, Yerlikaya, Yener, & Öktem, 2019) and the measure used was the resistance to interference in seconds as calculated by subtracting the time taken to read the colour names from the time taken to name the colour ink of written colour names. The Golden version of the Stroop test (Golden, 1978) was used in CR/RANN and number of words completed in 45 seconds on the Color-Word page was used as the measure. There was no comparable measure of response inhibition and selective attention in TILDA.

*Sustained Attention*

Sustained attention was assessed with the Psychomotor Vigilance Test (PVT; Dorrian, Rogers and Dinges, 2005) in CR/RANN using the number of false alarms (i.e. errors of commission) and the median reaction time across trials with an inter-trial interval of two to four seconds. It was assessed with the Sustained Attention to Response Task (SART; Robertson *et al.*, 1997) in TILDA using the number of errors of commission and the coefficient of variation in reaction time as measures. There was no comparable measure of sustained attention in DEU.

*Verbal Episodic Memory (Immediate)*

Immediate verbal episodic memory was assessed in DEU measure with the immediate recall score from the Öktem Verbal Memory Processes Test (OVMPT; Öktem, 1992) which is a validated Turkish version of the Rey Auditory Verbal Learning Test (RAVLT; Bosgelmez et al., 2015) and measures the number of words immediately recalled from a 15-item word list. The CR/RANN measure was the total recall score on the Selective Reminding Test (SRT; Buschke & Fuld, 1974), which measures the total number of words recalled from 6 trials of a 12-item word list (Strauss et al., 2006). The TILDA measure was the number of words immediately recalled from a 10-item word list.

##### *Verbal Episodic Memory (Delayed)*

Delayed verbal episodic memory was assessed in DEU using the delayed recall score from the OVMPT which consisted of the number of words recalled from the 15-word list after a 40 minute delay, and in CR/RANN using the delayed recall score from the SRT which consisted of the number of words recalled from the 12-item word list after an approximate 15 minute delay. It was assessed in TILDA using the number of words recalled after an approximate 20-25 minute delay from a 10-item word list (depending on length of time it took participants to complete intervening items).

##### *Verbal Episodic Memory (Learning)*

Verbal episodic memory learning was assessed in DEU using the OVMPT total learning score which was the total number of words recalled in each trial and in CR/RANN using the consistent long-term retrieval score on the SRT which was the number of words consistently recalled on all subsequent trials (Strauss et al., 2006). There was no comparable measure of verbal episodic memory learning in TILDA.

##### *Working Memory*

Working memory was assessed in DEU using both the Digit Span Forward and Digit Span Backward tests from the Wechsler Memory Scale – Revised Edition (WMS-R; Wechsler, 1987). In CR/RANN, the Letter-Number Sequencing test from the Wechsler Adult Intelligence Scale – Third Edition (WAIS-III; Wechsler, 1997) was used. There was no comparable measure of working memory in TILDA.

##### *Visuospatial Ability*

Visuospatial ability was assessed in DEU using the Judgement of Line Orientation Test (BLOT; Benton, Varney, & Hamsher, 1978) which measures participants' capacity to discriminate the direction of lines. In CR/RANN, the Block Design test from the WAIS-III, which measures ability of participants to replicate models or pictures presented to them using blocks (Strauss et al., 2006), was used. There was no comparable measure of visuospatial ability in TILDA.

#### Statistical Analysis

*Calculating significance of replications - Random Label Permutation*

For findings replicated in multiple datasets, the probability of obtaining p-values by chance was calculated by random-label permutation (Good, 1994) as follows:

1. BrainPAD scores were randomly shuffled using MATLAB's randperm.m.
2. Spearman's partial correlations were conducted between randomly shuffled brainPAD scores and the cognitive dependent variables, controlling for age and gender.
3. Steps 1- 2 were repeated one million times.
4. The number of instances in which random p-values were more extreme (i.e. smaller) than actual p-values were counted. This number was divided by one million to obtain the probability of this finding replicating across multiple datasets by chance. Replicated findings were deemed significant if this probability was less than .05.

The code used for this step is available here:
https://github.com/rorytboyle/brainPAD_dataAnalysis

*Multiple comparisons correction - Maximum Statistic Correction*

To control for the familywise error rate within individual datasets, a maximum statistic correction was applied as follows:

1. In each test set, brainPAD scores were randomly shuffled using MATLAB's randperm.m.
2. Spearman's partial correlations were conducted between the randomly shuffled brainPAD scores and the cognitive dependent variables, controlling for age and gender.
3. Steps 1-2 were repeated ten thousand times and the maximum rho value was stored each time.
4. Correlations between actual brainPAD scores and cognitive variables were deemed significant if they had a rho value greater than the rho value in the $95^{th}$ percentile of the maximum rho values.

This approach was a variant of max T adjustments (Dudoit, Shaffer, & Boldrick, 2003) and was used because Bonferroni and Šidák adjustments can be overly conservative when there are correlated dependent measures (Conneely & Boehnke, 2007; Dudoit et al., 2003). The code used to perform this correction is available here:
https://github.com/rorytboyle/brainPAD_dataAnalysis

***Table S.2***

*Cognitive measures available across each dataset in comparable cognitive domains (extended version of table 1 in main text).*

| **Cognitive Domain(s)** | **DEU Measure *(N)*** | **CR/RANN Measure *(N)*** | **TILDA Measure *(N)*** |
|---|---|---|---|
| **General Cognitive Status** | MMSE (total score) *(172)* | DRS (total score) *(370)* | MMSE (total score) *(485)* |
| **Premorbid Intelligence** | n/a | AMNART (total score) *(362)* | NART (total score) *(486)* |
| **Phonemic Verbal Fluency** | F-A-S Test (total number of words named for each letter) *(137)* | CFL Test (total number of words named for each letter) *(360)* | n/a |
| **Semantic Verbal Fluency** | Animals Test (total number of animals named) *(175)* | Animals Test (total number of animals named) *(361)* | Animals Test (total number of animals named) *(487)* |
| **Processing Speed** | TMT A (time to complete) *(93)* | TMT A (time to complete) *(361)* | CTT 1 (time to complete) *(487)* |
| **Processing Speed, Visual Attention, Cognitive Flexibility** | TMT B (time to complete) *(84)* | TMT B (time to complete) *(357)* | CTT 2 (time to complete) *(482)* |
| **Visual Attention, Cognitive Flexibility** | TMT B minus TMT A (difference in time to complete both tasks) *(84)* | TMT B minus TMT A (difference in time to complete both tasks) *(357)* | CTT 2 minus CTT 1 (difference in time to complete both tasks) *(482)* |
| **Cognitive Flexibility** | WCST Perseverative Error Percentage *(50)* | WCST Perseverative Error Percentage *(327)* | n/a |
| **Response Inhibition, Selective Attention** | Stroop (Turkish Capa version; (Emek-Savaş et al., 2019) Interference Score - Time (seconds taken to name colour ink of written colour names minus seconds taken to read colour names) *(150)* | Stroop (Golden version; Golden, 1978) Interference Score - Words (number of words completed in 45 seconds on Color-Word page) *(359)* | n/a |
| **Sustained Attention (Errors of Commission)** | n/a | PVT Number of False Alarms *(176)* | SART Number of Errors of Commission *(482)* |
| **Sustained Attention (Reaction Time)** | n/a | PVT Median Reaction Time (across trials with 2 to 4 second inter-trial interval) *(176)* | SART Coefficient of Variation in Reaction Time *(479)* |
| **Verbal Episodic Memory (Immediate)** | OVMPT Immediate Recall (number of words recalled from 15-item list) *(175)* | SRT Total Score (number of words recalled from 6 trials of 12-item list) *(360)* | Immediate Recall (number of words recalled from 10-item list) *(487)* |
| **Verbal Episodic Memory (Delayed)** | OVMPT Delayed Recall (number of words recalled from 15-item list after 40-minute delay) *(175)* | SRT Delayed Recall (number of words recalled from 15-item list after approximate 15-minute delay) *(360)* | Delayed Recall (number of words recalled from 10-item list after approximate 20-25 minute delay) *(487)* |
| **Verbal Episodic Memory (Learning)** | OVMPT Total Learning Score (total number of words recalled in each trial) *(175)* | SRT Consistent Long Term Retrieval (number of words consistently recalled on all subsequent trials) *(360)* | n/a |
| **Working Memory** | WMS-R Digit Span Forward Test *(171)* | WAIS-III Letter Number Sequencing Test *(360)* | n/a |
| | WMS-R Digit Span Backward Test *(170)* | | |
| **Visuospatial Ability** | BLOT *(80)* | WAIS-III Block Design Test *(356)* | n/a |

*Note: MMSE = Mini-mental state examination* (Folstein et al., 1975)*; DRS Total Score= Mattis Dementia Rating Scale-2 – Total Score* (Jurica et al., 2001)*; NART = National Adult Reading Test* (Nelson & Willinson, 1982)*; AMNART = American National Adult Reading Test* (Grober & Sliwinski, 1991)*; CTT = Colour Trails Test* (D'Elia et al., 1996)*; TMT = Trail Making Test (Reitan, 1955); WCST = Wisconsin Card Sorting Test* (Heaton et al., 1993)*; SART = Sustained Attention to Response Test* (Robertson et al., 1997)*; PVT = Psychomotor Vigilance Task* (Dorrian et al., 2005)*; OVMPT = Öktem Verbal Memory Processes Test (Öktem, 1992); SRT = Selective Reminding Test* (Buschke & Fuld, 1974)*; WMS-R = Wechsler Memory Scale (Wechsler, 1987); WAIS-III = Wechsler Adult Intelligence Scale – Third Edition (Wechsler, 1997); BLOT = Benton's Judgement of Line Orientation Test* (Benton et al., 1978)

### Supplementary Results

#### BrainPAD Model Performance

Application of the coefficients to all three test sets resulted in a correlation between brain predicted age and chronological age of r = 0.73 (p < 0.0001), which explained 52.79% of the variance ($R^2$). The combined test set had a mean chronological age of 62.75 (SD = 14.3) years at the time of scanning and mean brain-predicted age of 62.94 (SD = 13.3) years. Mean brainPAD was +0.18 (SD = 10.25) years. MAE was 8.33 years and RMSE was 10.24 years. Weighted MAE was 0.112 years.

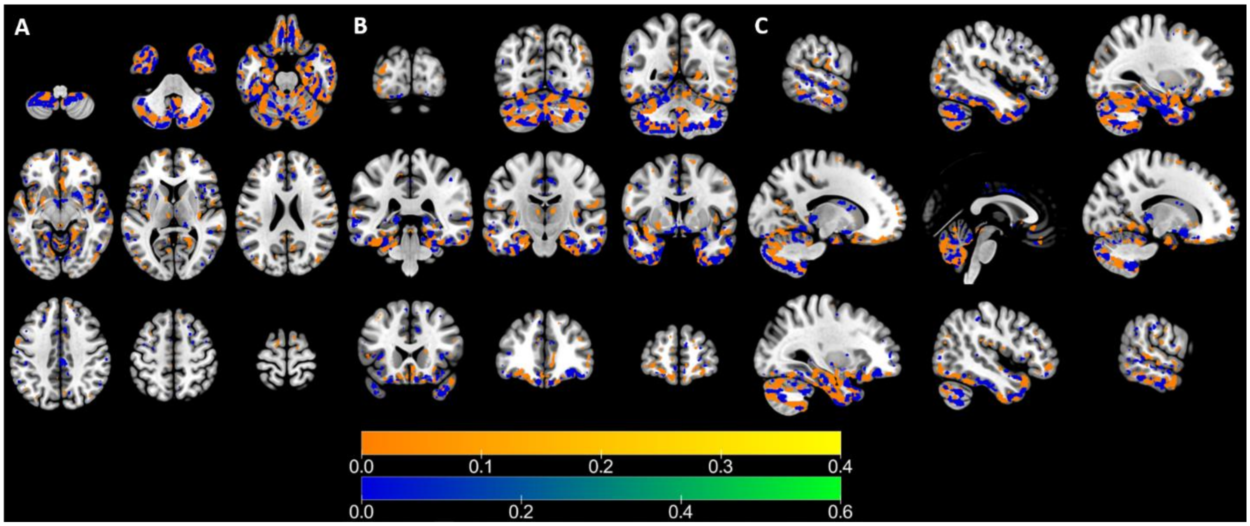


*Fig S.1: Voxels with negative coefficient values overlaid on voxels with positive coefficient values. A = Axial Plane, B = Coronal Plane, C = Sagittal Plane. Note: Signs of these voxels were flipped to aid plotting.*

#### Gender differences in brainPAD

Mean brainPAD differed significantly by gender in all datasets, Welch's t(1009.55) = - 5.79, $p < .0001$. Males (M = -1.81, SD = 9.92) had significantly lower brainPADs than females (M = 1.81, SD = 10.23) (see fig S.2). Within individual test sets, males had significantly lower brainPADs, compared to females, in in CR/RANN ($p < .0001$) and TILDA ($p < .0001$) but not in DEU ($p = 0.148$) (see fig S.3).

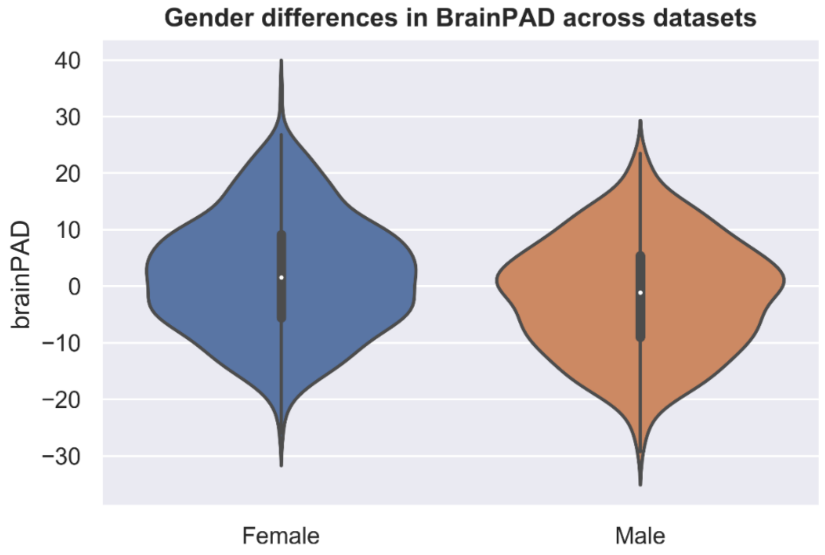


*Fig S.2. Violin plot comparing distributions of brainPADs between genders across all datasets*

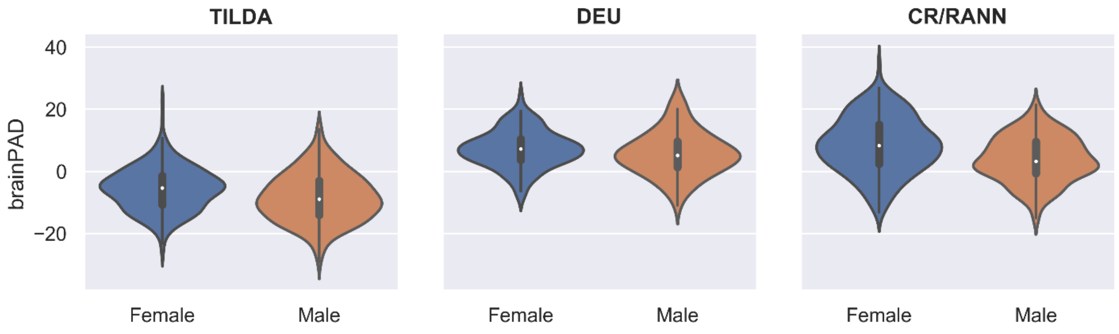


*Fig S.3: Violin plots comparing distributions of brainPADs between genders within datasets*

***Table S.3***

*Range, standard deviation (SD), and interquartile range (IQR) of variables used in partial correlations between brainPAD and cognitive functions.*

| Cognitive Domain | DEU | | | CR/RANN | | | TILDA | | |
|---|---|---|---|---|---|---|---|---|---|
| | Range | SD | IQR | Range | SD | IQR | Range | SD | IQR |
| **Age** | 45.95 | 8.59 | 13.08 | 61.00 | 17.09 | 30.25 | 38.00 | 7.21 | 8.00 |
| **BrainPAD** | 37.79 | 6.44 | 7.27 | 49.26 | 8.57 | 11.24 | 52.18 | 7.52 | 10.15 |
| **General Cognitive Status** | 16.00 | 2.98 | 4.00 | 16.00 | 2.85 | 4.00 | 9.00 | 1.41 | 2.00 |
| **Premorbid Intelligence** | n/a | | | 48.00 | 9.31 | 14.00 | 49.00 | 11.22 | 17.00 |
| **Phonemic Verbal Fluency** | 70.00 | 13.86 | 17.00 | 64.00 | 11.97 | 16.00 | | | |
| **Semantic Verbal Fluency** | 34.00 | 5.79 | 8.00 | 45.00 | 5.52 | 6.00 | 36.00 | 5.43 | 8.00 |
| **Processing Speed** | 267.00 | 34.79 | 33.00 | 93.45 | 10.93 | 12.00 | 211.88 | 24.28 | 27.35 |
| **Processing Speed, Visual Attention, Cognitive Flexibility** | 342.00 | 65.77 | 69.00 | 279.18 | 40.74 | 38.50 | 309.13 | 41.63 | 42.23 |
| **Visual Attention, Cognitive Flexibility** | 290.00 | 54.73 | 62.00 | 271.47 | 37.21 | 28.78 | 223.77 | 27.95 | 29.72 |
| **Cognitive Flexibility** | 40.12 | 7.68 | 7.99 | 66.67 | 8.69 | 8.18 | n/a | | |
| **Response Inhibition, Selective Attention** | 241.00 | 32.16 | 27.50 | 68.00 | 11.31 | 15.00 | n/a | | |
| **Sustained Attention (Errors of Commission)** | n/a | | | 8.00 | 1.50 | 2.00 | 23.00 | 3.71 | 4.00 |
| **Sustained Attention (Reaction Time)** | n/a | | | 572.00 | 75.97 | 74.50 | 1.30 | 0.16 | 0.15 |
| **Verbal Episodic Memory (Immediate)** | 11.00 | 1.97 | 3.00 | 47.00 | 9.79 | 14.00 | 10.00 | 1.48 | 2.00 |
| **Verbal Episodic Memory (Delayed)** | 15.00 | 4.51 | 7.00 | 19.00 | 2.48 | 4.00 | 10.00 | 2.46 | 3.00 |
| **Verbal Episodic Memory (Learning)** | 67.00 | 29.33 | 45.00 | 71.00 | 17.54 | 26.25 | n/a | | |
| **Working Memory** | 5.00† | 1.21 | 1.00 | 16.00 | 3.13 | 4.00 | n/a | | |
| | 7.00‡ | 1.17 | 2.00 | n/a | | | n/a | | |
| **Visuospatial Ability** | 19.00 | 4.45 | 6.00 | 60.00 | 13.22 | 20.25 | n/a | | |

*Note: † Digit Span Forwards; ‡ Digit Span Backwards*

#### BrainPAD and Cognitive Function – Replicated Findings

##### *General Cognitive Function*

BrainPAD was significantly negatively correlated with measures of general cognitive status in both DEU (MMSE) and CR/RANN (DRS) but not TILDA (MMSE). The probability of finding p-values as extreme as these actual p-values in the three datasets is indicated by the replication p-value, which was 0.000002.

##### *Semantic Verbal Fluency*

BrainPAD was significantly negatively correlated with semantic verbal fluency as measured by the Animals test in DEU and CR/RANN, but not TILDA, replication p-value < 0.000001.

##### *Processing Speed, Visual Attention, and Cognitive Flexibility*

BrainPAD was significantly positively correlated with measures of visual attention and cognitive flexibility in DEU (TMT B), CR/RANN (TMT), and TILDA (CTT 2), replication p-value = 0.000054.

##### *Visual Attention and Cognitive Flexibility*

BrainPAD was significantly positively correlated with measures of visual attention and cognitive flexibility in DEU (TMT B – TMT A) and CR/RANN (TMT B – TMT A), but not in TILDA (CTT 2 – CTT 1), replication p-value = 0.000966.

#### BrainPAD and Cognitive Function – Non-replicated Findings

Additionally, higher brainPAD scores were also significantly correlated with reduced performance on other measures of cognitive function but these correlations were not replicated in another dataset.

##### *Phonemic Verbal Fluency*

BrainPAD was significantly negatively correlated with phonemic verbal fluency as measured by the KAS test in DEU (rho = -0.326, p = 0.0001) and this survived multiple comparison correction. However, brainPAD was not significantly correlated with the CFL test performance in CR/RANN (rho = -0.0771, p = 0.1454). There was no comparable measure in TILDA.

##### *Premorbid Intelligence*

BrainPAD was significantly negatively correlated with premorbid intelligence as measured by the AMNART in CR/RANN (rho = -0.2322, p < 0.00001) and this survived multiple comparison correction. However, brainPAD was not significantly correlated with the NART in TILDA (rho = -0.0485, p = 0.2873). There was no comparable measure in DEU.

##### *Verbal Episodic Memory (Learning)*

BrainPAD was significantly negatively correlated with verbal episodic memory learning as measured by the OVMPT Total Learning Score in DEU (rho = -0.3196, p < 0.00001) and this survived multiple comparison correction. However, brainPAD was not significantly correlated with the consistent long-term retrieval score on the SRT in CR/RANN (rho = 0.0657, p = 0.2151). There was no comparable measure in TILDA.

##### *Visuospatial Ability*

BrainPAD was significantly negatively correlated with visuospatial ability as measured by the Block Design test in CR/RANN (rho = -0.1824, p = 0.0006) and this survived multiple comparison correction. However, brainPAD was not significantly correlated with the BLOT in DEU (rho = -0.0809, p = 0.4815). There was no comparable measure of visuospatial ability in TILDA.

#### BrainPAD and Cognitive Function – Non-significant Findings

##### *Processing Speed*

BrainPAD was significantly positively correlated with processing speed as measured by the CTT 1 in TILDA (rho = 0.1208, p = 0.0077) but this result did not survive multiple comparison correction. Furthermore, brainPAD was not significantly correlated with the TMT A in DEU (rho = 0.1232, p = 0.2448) nor in CR/RANN (rho = 0.0595, p = 0.2610).

##### *Response Inhibition and Selective Attention*

BrainPAD was significantly correlated with reduced response inhibition and selective attention as measured by the number of incongruous words completed in 45 seconds on the Stroop Color-Word test in CR/RANN (rho = -0.1755, p = 0.0009). However, this result did not survive multiple comparisons correction. Furthermore, brainPAD was not significantly correlated with the Stroop Color-Word test in DEU (rho = 0.0854, p = 0.302).

##### *Working Memory*

BrainPAD was significantly negatively correlated with working memory as measured by the WMS-R Digit Span Backward test in DEU (rho = -0.2974, p = 0.0001) but this result did not survive multiple comparisons correction. Furthermore, brainPAD was not significantly correlated with the WMS-R Digit Span Forward test in DEU (rho = -0.1310, p = 0.0895) or the WAIS-III Letter-Number Sequencing test in CR/RANN (rho = -0.0469, p = 0.3759).

##### *Verbal Episodic Memory (Immediate)*

BrainPAD was significantly negatively correlated with immediate verbal episodic memory as measured by the OVMPT Immediate Recall score in DEU (rho = -0.2194, p = 0.0037) but this result was not significant after correction for multiple comparisons. Moreover, brainPAD was not significantly correlated with immediate recall in TILDA (rho -0.0374, p = 0.4114) or total recall score on the Selective Reminding Test in CR/RANN (rho = 0.0407, p = 0.4428).

##### *Verbal Episodic Memory (Delayed)*

BrainPAD was significantly negatively correlated with delayed verbal episodic memory as measured by the OVMPT Delayed Recall score in DEU (rho = -0.2797, p = 0.0002) but this result was not significant after correction for multiple comparisons. Additionally, brainPAD was not significantly correlated with delayed recall in TILDA (rho = 0.0122, p = 0.7887) or delayed recall on the Selective Reminding Test in CR/RANN (rho = 0.0343, p = 0.5173).

##### *Sustained Attention*

BrainPAD was not significantly correlated with the number of commission errors on the SART (rho = 0.0499, p = 0.2752) in the TILDA dataset nor on the PVT (rho = 0.0203, p = 0.7902) in CR/RANN. Moreover, brainPAD was not significantly correlated with response variability, as measured by the coefficient of variation in reaction times, in the SART in TILDA (rho = 0.0436, p = 0.3425) nor in the PVT in CR/RANN (rho = -0.0212, p = 0.7813).

##### *Cognitive Flexibility*

BrainPAD was not significantly correlated with percentage of perseverative errors on the WCST in the DEU dataset (rho = 0.0722, p = 0.6258) and in CR/RANN (rho = 0.0429, p = 0.4411).

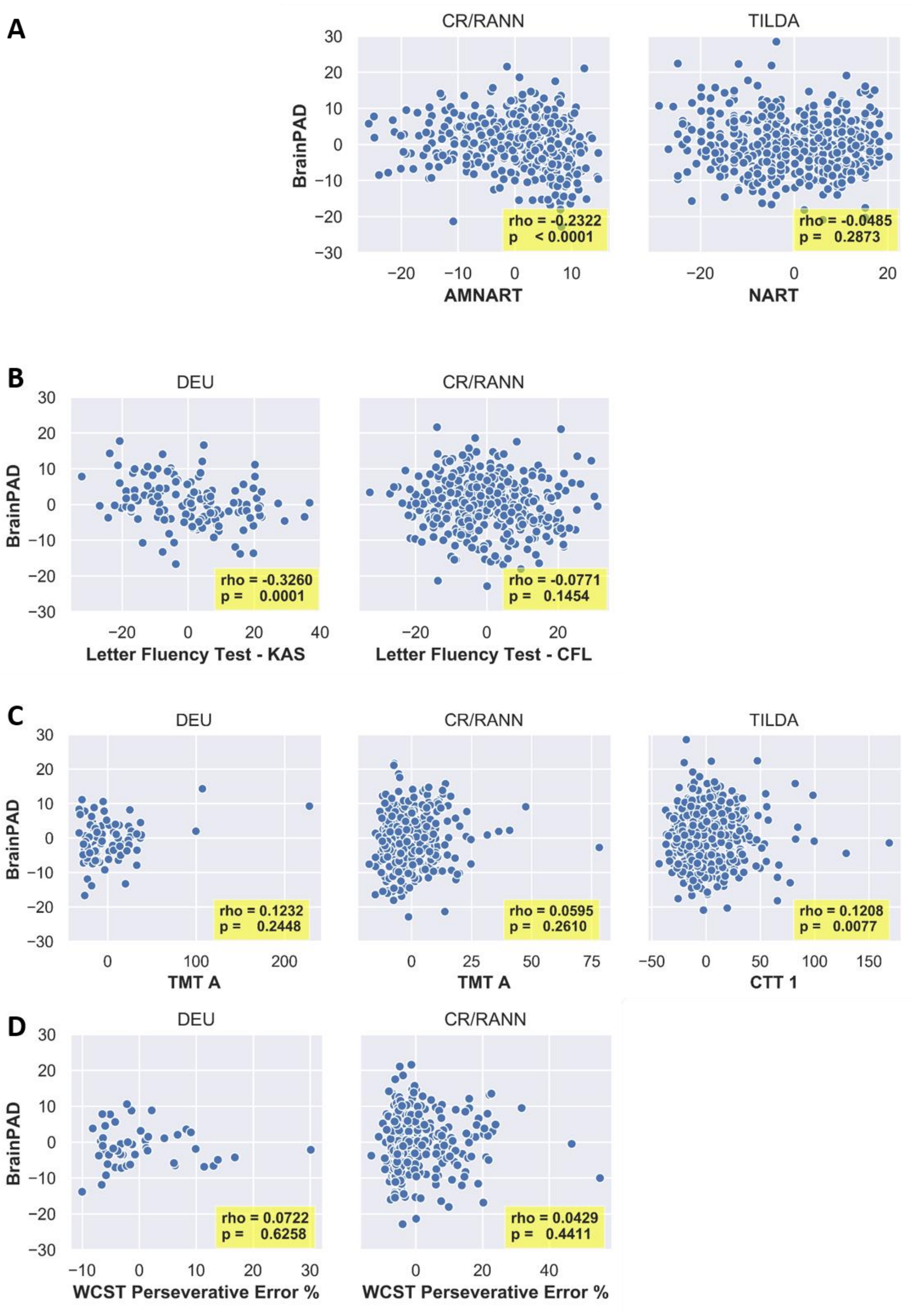
A
CR/RANN
TILDA
BrainPAD
rho = -0.2322
p < 0.0001
rho = -0.0485
p = 0.2873
AMNART
NART
B
DEU
CR/RANN
rho = -0.3260
p = 0.0001
rho = -0.0771
p = 0.1454
Letter Fluency Test - KAS
Letter Fluency Test - CFL
C
DEU
CR/RANN
TILDA
rho = 0.1232
p = 0.2448
rho = 0.0595
p = 0.2610
rho = 0.1208
p = 0.0077
TMT A
TMT A
CTT 1
D
DEU
CR/RANN
rho = 0.0722
p = 0.6258
rho = 0.0429
p = 0.4411
WCST Perseverative Error %
WCST Perseverative Error %

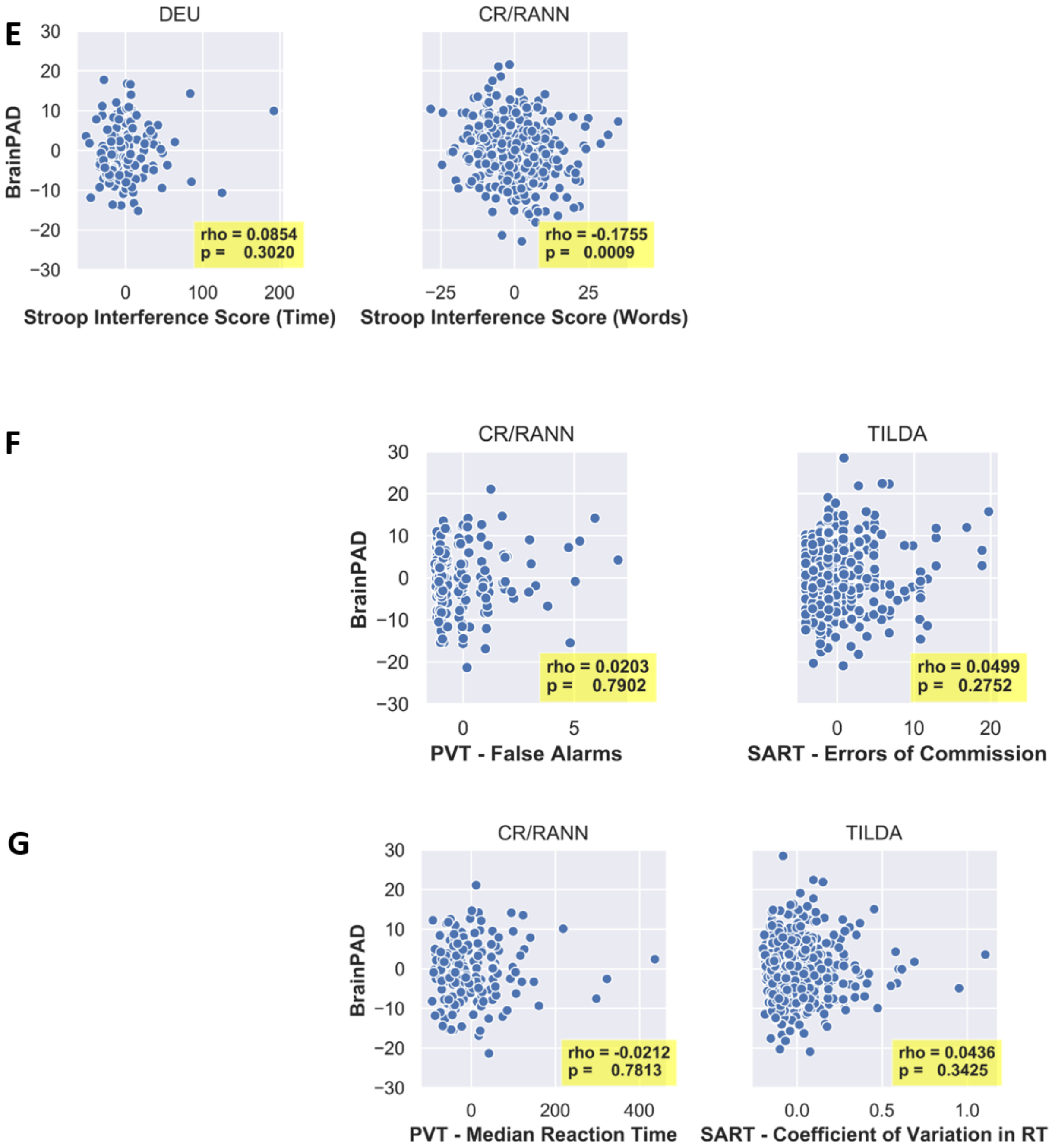
E
DEU
CR/RANN
BrainPAD
rho = 0.0854
p = 0.3020
rho = -0.1755
p = 0.0009
Stroop Interference Score (Time)
Stroop Interference Score (Words)
F
CR/RANN
TILDA
BrainPAD
rho = 0.0203
p = 0.7902
rho = 0.0499
p = 0.2752
PVT - False Alarms
SART - Errors of Commission
G
CR/RANN
TILDA
BrainPAD
rho = -0.0212
p = 0.7813
rho = 0.0436
p = 0.3425
PVT - Median Reaction Time
SART - Coefficient of Variation in RT

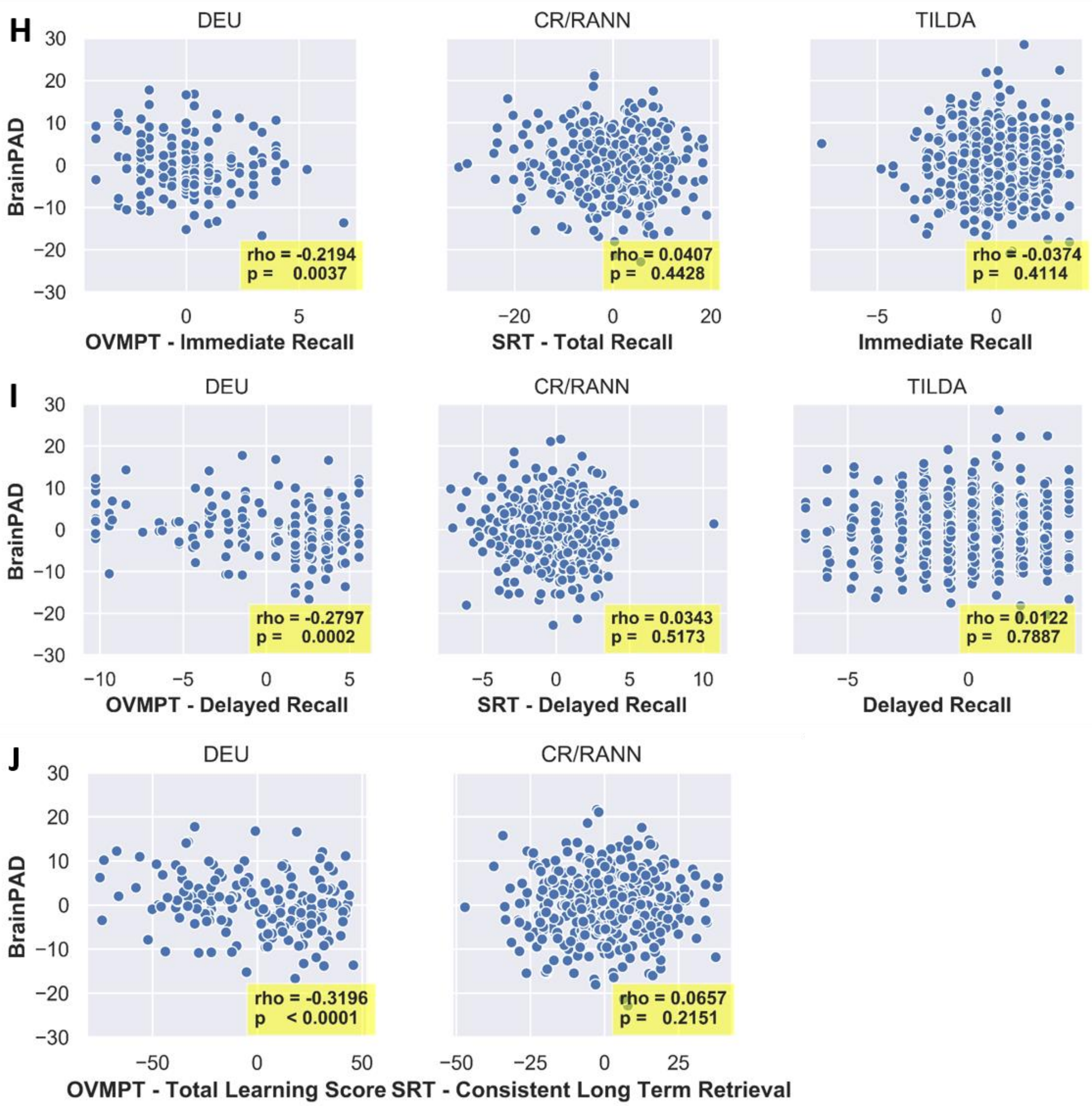

H
DEU
CR/RANN
TILDA
BrainPAD
OVMPT - Immediate Recall
SRT - Total Recall
Immediate Recall
rho = -0.2194
p = 0.0037
rho = 0.0407
p = 0.4428
rho = -0.0374
p = 0.4114
I
DEU
CR/RANN
TILDA
BrainPAD
OVMPT - Delayed Recall
SRT - Delayed Recall
Delayed Recall
rho = -0.2797
p = 0.0002
rho = 0.0343
p = 0.5173
rho = 0.0122
p = 0.7887
J
DEU
CR/RANN
BrainPAD
OVMPT - Total Learning Score
SRT - Consistent Long Term Retrieval
rho = -0.3196
p < 0.0001
rho = 0.0657
p = 0.2151

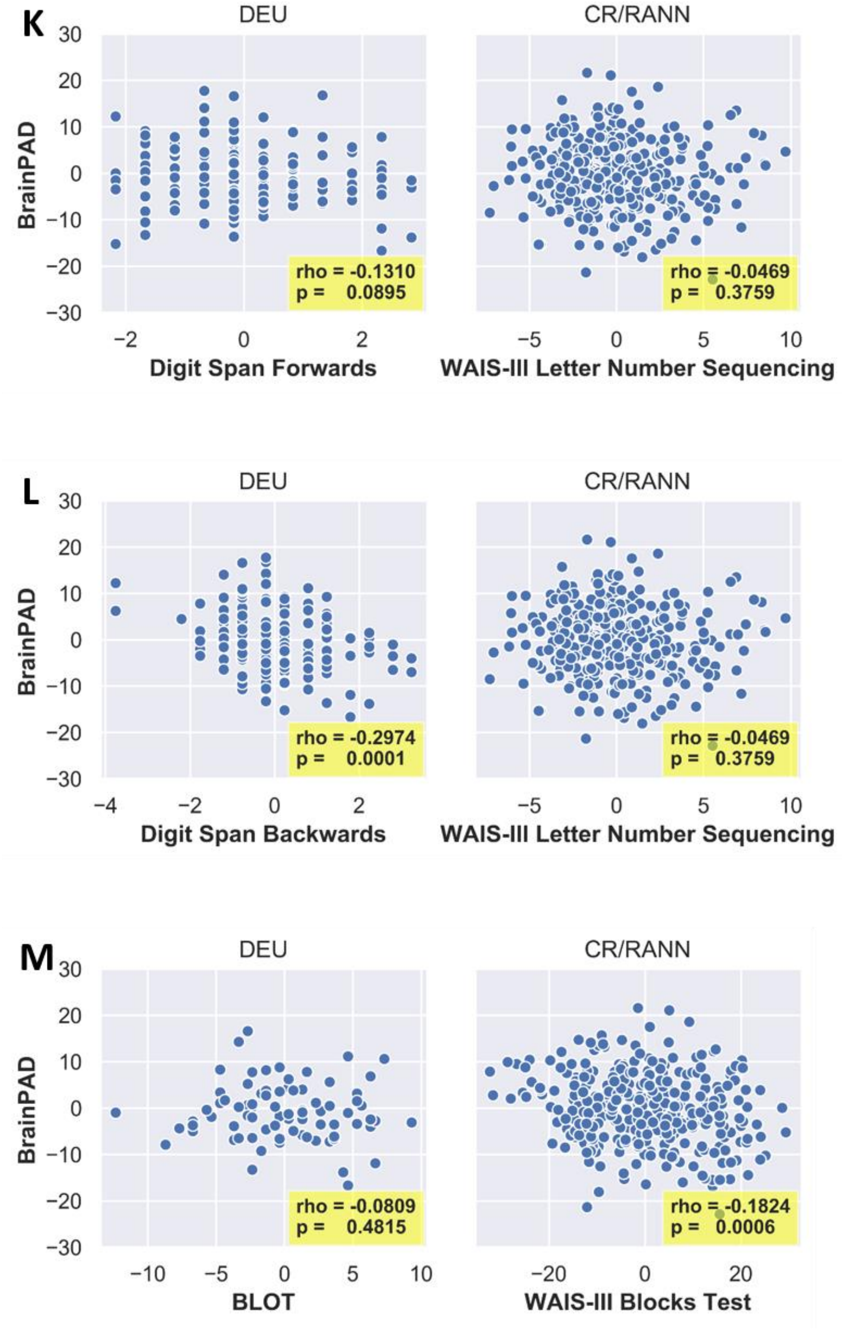


*Fig S.4: Scatterplots of non-replicated correlations between the residuals of brainPAD and cognitive measures after regressing brainPAD on age and gender, and regressing each cognitive measure on age and gender. A: Premorbid Intelligence; B: Phonemic Verbal Fluency; C: Processing Speed; D: Cognitive Flexibility; E: Response Inhibition and Selective Attention; F: Sustained Attention (Errors of Commission); G: Sustained Attention (Reaction Time); H: Verbal Episodic Memory (Immediate); I: Verbal Episodic Memory (Delayed); J: Verbal Episodic Memory (Learning); K: Working Memory; L: Working Memory; M: Visuospatial Ability.*
